# Human monoclonal antibodies that target the SFTSV glycoprotein Gn head from four neutralizing epitope groups

**DOI:** 10.1101/2025.11.26.690690

**Authors:** Qianran Wang, Hao Li, Fanchong Jian, Aoxiang Han, Yuanni Liu, Jingyi Liu, Yuanling Yu, Jing Wang, Lingling Yu, Yanxia Wang, Haiyan Sun, Miaomiao Ma, Fei Shao, Liuluan Zhu, Wei Liu, Yunlong Cao

**Affiliations:** Changping Laboratory, Beijing, P.R. China; State Key Laboratory of Pathogen and Biosecurity, Academy of Military Medical Sciences, Beijing, P.R. China; Biomedical Pioneering Innovation Center (BIOPIC), School of Life Sciences, Peking University, Beijing, China; Department of Infectious Diseases, Yantai Qishan Hospital, Yantai, P.R. China; College of Future Technology, Peking University, Beijing, P. R. China; Beijing Key Laboratory of Viral Infectious Diseases, Beijing Ditan Hospital, Capital Medical University, Beijing, P.R. China; Peking-Tsinghua Center for Life Sciences, Peking University, Beijing, P.R. China

**Author notes:** These authors contributed equally (Qianran Wang, Hao Li). Correspondence (Y.C.), (W.L.), (L.Z.).

## Abstract

Severe fever with thrombocytopenia syndrome virus (SFTSV) is a lethal bunyavirus lacking approved countermeasures. From SFTS survivors, we isolate 84 human monoclonal antibodies (mAbs) against the viral glycoproteins Gn and Gc. Gn-specific mAbs demonstrate superior neutralization breadth and potency compared to the restricted neutralizing activity observed with Gc. Using a high-throughput yeast display deep mutational scanning (DMS) platform, we classify Gn-head mAbs into eight epitope groups, among which four groups (IA, ID, IIIA, IIIB) conferring neutralization. Notably, mAbs BD70-4003 (group IA) and BD70-4017 (group IIIA) exhibit broad neutralization and provide 100% protection in a lethal mouse model. Cryo-EM structural analysis of these mAbs in complex with the Gn-head reveal their binding interfaces, directly validating the epitope residues identified by DMS. Our study delineates the antigenic landscape of SFTSV Gn, identifies potent therapeutic candidates, and establishes DMS coupled with structural validation as a powerful framework for antibody discovery against bunyavirus.

## INTRODUCTION

Severe fever with thrombocytopenia syndrome virus (SFTSV), or *Bandavirus dabieense*, is an emerging tick-borne phlebovirus first identified in China in 2009 and has since reported across East and Southeast Asia, including Japan, South Korea, Vietnam, Pakistan, and Thailand.^1–3^ Clinical manifestations range from fever, thrombocytopenia, and leukopenia to multi-organ dysfunction, with a subset of patients developing hemorrhagic symptoms and organ failure. The case fatality rate (CFR) is approximately 16.2% (95% CI 14.6-17.8) among confirmed cases.^4–6^ The principal vector, the Asian longhorned tick, has recently been detected in North America, Russia, and Australia, raising concerns regarding geographic expansion and the increasing risk of human-to-human transmission.^7–9^ Despite its designation by the World Health Organization as a priority pathogen requiring urgent countermeasures, no licensed vaccines or effective antiviral drugs are currently available.

The tripartite SFTSV genome encodes a glycoprotein precursor on the M segment, which is cleaved into Gn and Gc subunits that mediate receptor binding and membrane fusion during viral entry.^10,11^ These glycoproteins are the primary mediators of viral entry; Gn is generally implicated in initial receptor attachment, while Gc facilitates the low-pH-dependent fusion of the viral and endosomal membranes. These Gn/Gc heterodimers organize into higher-order hexon and penton structures, creating a dense and ordered lattice that covers the viral envelope.^12^

Recent structural biology breakthroughs have suggested that the Gn glycoprotein as an immunodominant target for neutralizing antibodies (mAbs).^12^ The Gn ectodomain consists of a globular head and a stem region. The Gn-head (residues 20-340) is further subdivided into three distinct structural domains: Domain I, Domain II, and Domain III. Domain I is characterized by high surface exposure and is believed to harbor the critical interfaces for host receptor engagement. Domain II acts as a β-connector structure, while Domain III forms a structural “cap” that partially shields the fusion loop of the Gc subunit, thereby regulating the timing of membrane fusion. The preservation of these domains across diverse SFTSV clades makes them attractive targets for the development of universal therapeutics and vaccine antigens.

Multiple monoclonal antibodies (mAbs) against Gn have been described, but their epitopes, potency, and breadth vary significantly.^13–19^ A systematic evaluation is essential to delineate the precise relationship between Gn-directed antibody epitopes and neutralization potency, both *in vitro* and *in vivo*, to inform the development of therapeutics and vaccines. Deep mutational scanning (DMS) technology enables epitope mapping with significantly higher efficiency and resolution than traditional binning techniques.^20,21^ We previously developed a high-throughput epitope mapping platform based on DMS to study SARS-CoV-2, establishing a pipeline to isolate large panels of mAbs and simultaneously identify their precise binding epitopes.^22–27^ Here, we applied this technology to SFTSV to elucidate the intrinsic relationship between antigenic epitopes and functional antibody profiles.

We report the isolation and epitope classification of human mAbs from SFTSV convalescents. We assessed binding and neutralization using enzyme-linked immunosorbent assay (ELISA), pseudovirus assays, and authentic virus neutralization assays. Using a novel yeast DMS approach adapted for bunyaviruses, we classified Gn-head mAbs into eight epitope groups (ⅠA-ⅠD, ⅡA-ⅡB, ⅢA-ⅢB). Four groups (ⅠA, ID, ⅢA and ⅢB) demonstrated neutralizing activity. Two antibodies in particular—BD70-4003 (IA) and BD70-4017 (IIIA)—exhibited broad neutralization and provided 100% protection in mice, making them promising drug candidates. Cryo-electron microscopy (cryo-EM) structural analyses of three mAbs (BD70-4003, BD70-4008 and BD70-4017) defined the molecular basis of recognition and verified the accuracy of our DMS system. Our results elucidate the overall epitope distribution of the SFTSV Gn-head domain and identify highly potent neutralizing antibodies targeting distinct epitopes, providing theoretical guidance for future therapeutic and vaccine design.

## RESULTS

### Isolation of mAbs from SFTS convalescents

We collected whole-blood samples from 12 SFTSV survivors who had been infected for more than one year (Table S1). Serum samples were analyzed for viral Gn/Gc-specific IgG titers and for neutralizing activities against SFTSV pseudovirus. The results showed that Gn or Gc reactive antibody remained at high levels in these serum samples (Figure 1A), and the serum exhibited high neutralizing activity against six pseudovirus strains (Figure 1B). This is consistent with previous reports that SFTSV glycoproteins induce long-lasting antibodies, and also suggests the possibility of potentially broadly neutralizing antibodies.^28,29^

**Figure 1.**
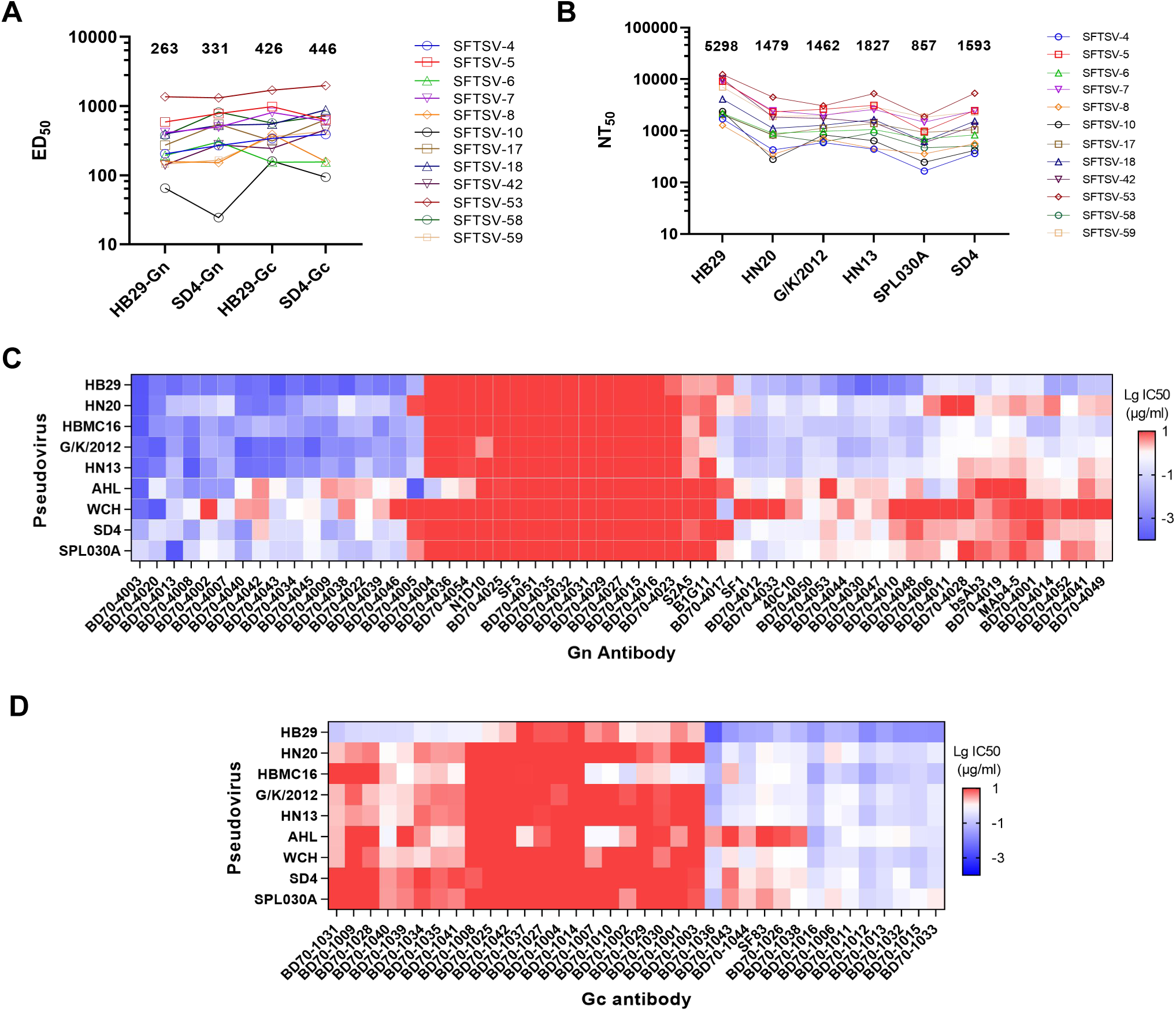
Identification of broad neutralizing antibodies from SFTSV-infected convalescents. (A) The median effective dose (ED_50_) of plasma from SFTSV-infected convalescents was determined by ELISA. The average value of each group was marked. (B) The half maximal neutralization titre (NT_50_) of plasma against VSV-based SFTSV pseudoviruses. (C-D) Neutralizing median inhibition concentration (IC_50_, μg/ml) of SFTSV Gn (C) or Gc (D) antibodies using VSV-based pseudoviruses. Neutralization assays were conducted in at least two biological replicates. See also Figure S1 and Tables S1-S5.

To isolate monoclonal antibodies, we purified antigen-specific memory B cells from peripheral blood mononuclear cells (PBMCs) using fluorescence-activated cell sorting (FACS). From the PBMCs, CD19⁺ IgM⁻ CD27⁺ memory B cells were stained simultaneously with HB29 Gn and Gc glycoproteins, and cells positive for either antigen were sorted as antigen-specific B cells (Figure S1A). To distinguish Gn- and Gc-specific responses during sequencing, each antigen was labeled with a unique DNA barcode. In total, we retrieved 1,316 paired heavy-light chain antigen-specific antibody sequences via high-throughput single-cell V(D)J sequencing.

A subset of antibody sequences was randomly selected for expression and further functional characterization. In total, we expressed 54 Gn-specific and 44 Gc-specific IgG1 antibodies *in vitro*. ELISA confirmed binding activity in 49/54 (90.7%) of the Gn group and 35/44 (79.5%) of the Gc group (Table S2). ELISA-positive Gn (n = 49) and Gc (n = 35) mAbs were subsequently subjected to neutralization assays.

In the neutralization assay, we employed 9 vesicular stomatitis virus (VSV)-based pseudovirus strains representing six major SFTSV genotypes (A–F) (Table S3-S6).^30–32^ These strains include: HB29 (genotype D), HBMC16 (genotype D), WCH (genotype A), HN13 (genotype A), HN20 (genotype F), G/K/2012 (genotype F), SD4 (genotype E), SPL030A (genotype B), and AHL (genotype C). The results show that mAbs against Gn exhibited more potent neutralization effects against the pseudovirus than those against Gc, which is consistent with previous reports that neutralizing antibodies against SFTSV primarily target the Gn antigen (Figures 1C and 1D).^14,33–35^ This observation was further validated in authentic virus neutralization assays, where eleven selected Gn-specific pseudovirus neutralizing mAbs—but zero Gc-specific mAbs—showed greater than 50% inhibition against the HBMC strain at 0.25 μg/mL (Figure S1B). Based on these results, we confirm that Gn is the immunodominant antigen target for SFTSV neutralizing antibodies.

### Epitope mapping via deep mutational scanning

Defining antibody epitopes is essential for understanding neutralization mechanisms and for identifying functionally important regions on viral antigens that can be targeted for therapeutic or vaccine development. To provide a high-resolution epitope map of the Gn antigenic surface, a comprehensive deep mutational scanning (DMS) platform was implemented.^21,36–38^ A yeast display library of the HB29 Gn-head domain (residues 20-340) was constructed, incorporating nearly all possible amino acid substitutions at each position. By subjecting this library to antibody selection and high-throughput sequencing of escape variants, critical binding residues were identified for 44 newly isolated human Gn-mAbs and several benchmark control antibodies. Specially, we performed negative selection FACS to isolate yeast cells that failed to bind specific antibodies, followed by sequencing to identify critical escape mutants. Unsupervised clustering of the resulting escape profiles partitioned the Gn-head into eight distinct epitope groups, providing a systematic framework for understanding antibody function.

The eight epitope groups (IA-ID, IIA-IIB, IIIA-IIIB) demonstrate a diverse distribution across the Gn-head structure (Figures 2A-C, Figure S2). Groups IA, IC, and ID are localized to Domain I. Group IA, represented by elite neutralizing mAbs such as BD70-4003, targets a surface-exposed region involving residues D159, G161, S163-S164, and L166. Group IC (e.g., control antibody SF5^39^) and ID (e.g., control antibody 40C10^16^) target distinct sites on the opposite faces of Domain I. Domain II is recognized by groups IIA and IIB, which include antibodies like N1D10 and B1G11^15^. No convergent V(D)J gene usage was observed within mAbs targeting Domain I and II (Figures 2D). Despite high binding scores, antibodies targeting Domain II often exhibit poor neutralizing activity, likely due to the domain’s less accessible orientation on the mature virion (Figures 2E and 2F).

**Figure 2.**
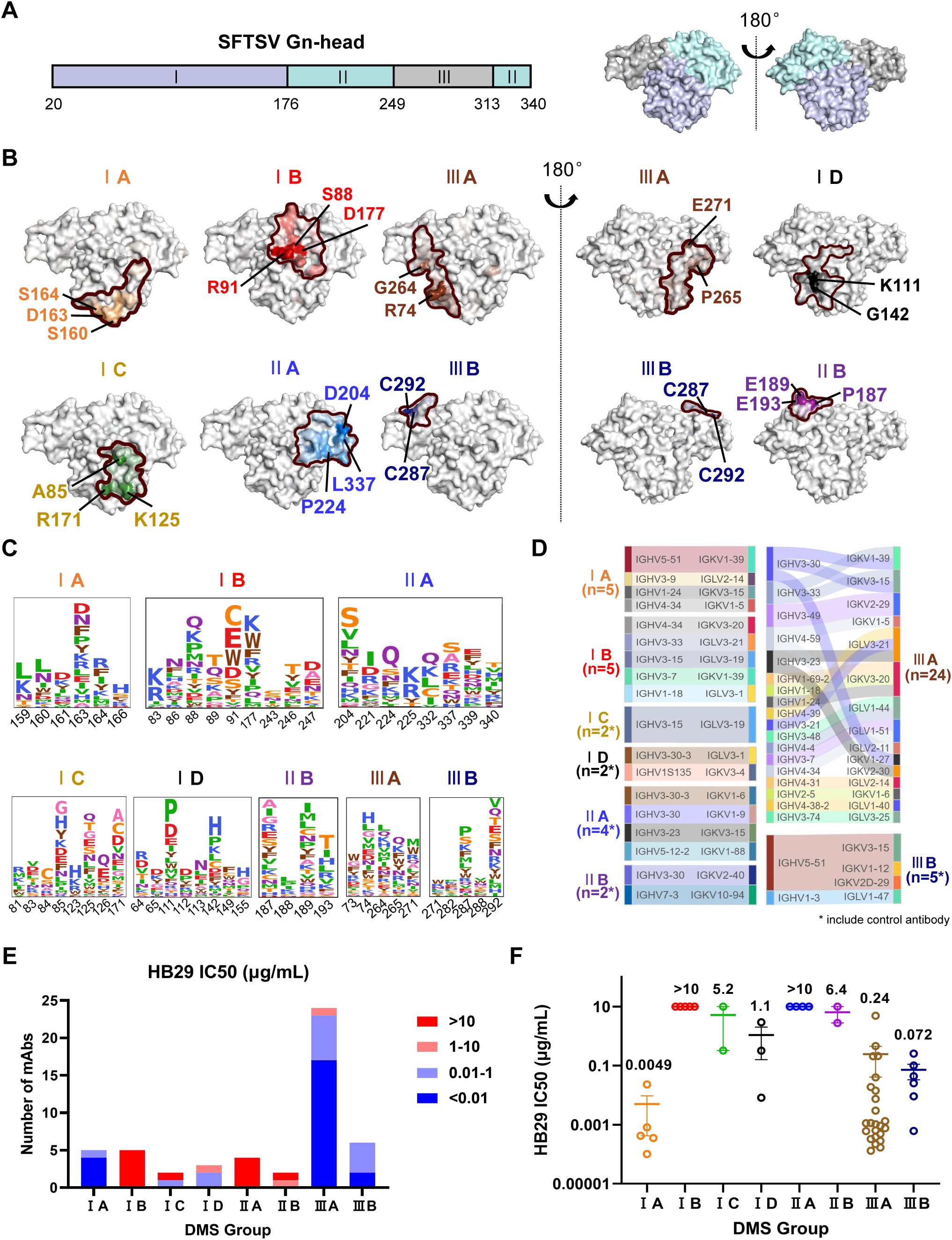
Epitope analysis of all Gn antibodies by yeast DMS. (A) Structural schematic of SFTSV Gn-head. The left panel illustrates the delineation of the three domains, colored in blue-white (Ⅰ), pale cyan (Ⅱ), and gray (Ⅲ), respectively; the corresponding structural schematic is shown in the right panel (PDB: 5Y10). (B) Structural projections and key epitope annotations of the eight DMS groups with different colors. The region outlined by the curved line corresponds to the epitope of each respective group. In the structural projection, amino acid residues with higher DMS scores are displayed in darker colors. (C) Average escape scores of antibodies in eight groups. Colors are assigned according to the type of amino acids, and the heights of the letters in the logo plot indicate escape scores. Residues are numbered based on Gn-head numbering. (D) The gene analysis of the antibodies in eight groups. Sankey diagram was generated to visualize the correspondence between heavy and light chains for the antibody genes in each group, with asterisks indicating the inclusion of control antibodies. (E-F) Statistical analysis of the number (E) and IC_50_ values (mean SEM) (F) for neutralization against the HB29 strain across DMS-classified antibody groups was performed. See also Figures S2-S3 and Tables S6-S7.

Group III antibodies are subdivided into IIIA and IIIB and target Domain III. Group IIIA antibodies, such as BD70-4017, interact with a cluster of residues including Q73-R74, G264-P265, and E271. Group IIIB antibodies, including the well-characterized MAb4-5^13^, focus on an epitope involving residues E271, G282, C287, K288, and C292. Notably, gene analysis of Group IIIB mAbs showed frequent genes involving IGHV5-51 (Figures 2D). Importantly, comparative analysis reveals that the most potent neutralizing activity is concentrated in Group IA and IIIA, followed by IIIB and ID, whereas IC, IIA, and IIB are largely non-neutralizing. This trend was also consistent across another eight SFTSV pseudoviruses strains, with the most potent neutralization again observed in the IA and IIIA antibody groups, highlighting the functional dominance of the IA and IIIA epitopes (Figure S3).

To validate the accuracy of our DMS platform, we compared our results with previously published crystal or cryo-EM structures. It has been reported that the interface between the Gn head and MAb4-5 (ⅢB) was located on sites S251-F256, K275-G278, and 284-294 by crystal structure analysis,^13^ which was consistent with the key amino acids (C287-K288, C292) we have screened (Figures 2B and 2C). For domain I antibodies SF5 (ⅠC) and 40C10 (ⅠD), DMS identified key residues S81-A85, S123-G126, R171 for SF5, and H64-S65, K111-K113, G142 for 40C10 (Figures 2B and 2C). Actually, SF5 (ⅠC) bound to sites R74-S90, G121-G126, T150-R171^39^, while 40C10 (ⅠD) bound to sites R62-Q66, D102, K111-S115, C156.^16^ Except for site G142, which was close to S65, all sites in this study were consistent with their complex structures. Results of mAbs against domain Ⅱ were also relatively accurate. According to Ren *et al*’s study,^15^ N1D10 (ⅡA) and B1G11 (ⅡB) mainly bound to sites E114-G121, G218-D226, V339-N340, and P185-E193, T321-V323, respectively, which also include the sites of our DMS results (Figures 2B and 2C). These correlations confirm that the yeast-based DMS library effectively captures the functional epitopes of SFTSV glycoproteins, providing a reliable high-throughput epitope determination alternative to structural mapping.

### Protective effect of Gn mAbs *in vitro* and *in vivo*

Previous pseudovirus and authentic virus neutralization screening identified six Gn antibodies (BD70-4003, BD70-4008, BD70-4009, BD70-4013, BD70-4017, and BD70-4022) with broad inhibitory effects (Table S3). These elite candidates were also found to have undergone extensive affinity maturation, as evidenced by high somatic hypermutation (SHM) rates and binding affinities (Table S7 and Figure S4). Notably, the binding affinity of BD70-4008 and BD70-4003 against Gn have reached in the low picomolar range.

Focus reduction neutralization test (FRNT) was performed to further validate the neutralization capacity of these elite mAbs (Figure 3A).^18^ All 6 Gn mAbs demonstrated potent neutralization *in vitro* with low FRNT_50_ values (Figure 3B). Interestingly, although BD70-4017 showed relatively modest neutralization in the pseudovirus system compared to its peers, its performance against authentic SFTSV strains was dramatically enhanced, illustrating that pseudotype assays may not fully recapitulate the inhibitory potential of certain epitopes.

**Figure 3.**
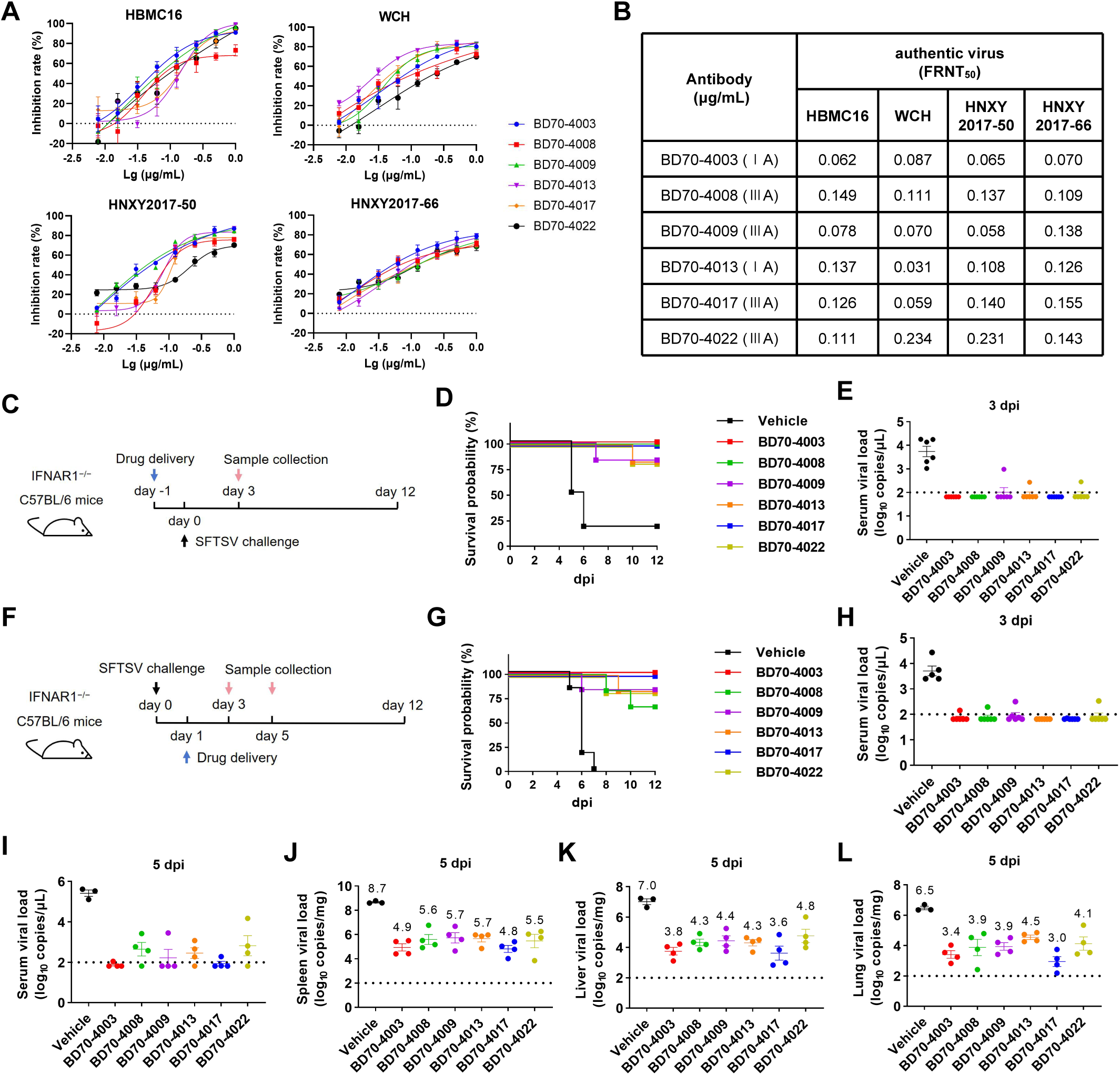
MAb-mediated protection of authentic virus challenge *in vitro* and *in vivo*. (A-B) The neutralization activity of six selected mAbs against four SFTSV authentic virus. The FRNT_50_ was calculated using the equation of log [inhibitor] versus response-variable slope (four parameters) method. Data were shown as mean SD values from three biologically independent replicates. (C and F) Schematic of the design for the HBMC16 challenge experiment in mice. In prophylactic groups (C), 5 mg/kg mAb was administered intraperitoneally and challenged by HBMC16 intraperitoneally 1 day later (C). In therapeutic groups (F), antibodies were given intraperitoneally 1 day after virus challenge. Each group consists of 6 mice. (D and G) Survival curves of mice in each group. (E, H and I) Viral load in the serum of mice in each group. (J-L) Viral load (mean SD) of mice challenged by HBMC16 in spleen (J), liver (K), or lung (L). Each point corresponds to samples from a mouse. The limit of detection is about 1.81×10^2^ copies/g and is shown as the dashed line. See also Figures S4.

To evaluate the prophylactic and therapeutic efficacy of the 6 mAb candidates against SFTSV *in vivo*, *IFNAR1^-/-^* mice were challenged with SFTSV before or after treatment with the mAb. In prophylactic and therapeutic groups, 5 mg/kg antibody was given to the mice via intraperitoneal (i.p.) injection 1 day pre-infection or 1 day post infection (dpi), respectively (Figures 3C and 3F). A vehicle group of mice that received a human IgG1 isotype control antibody was also included. Each group consisted of five or six mice. The body weight of each mouse in each group was monitored and recorded daily. Serum samples were collected from all mice on 3 dpi, with an additional collection performed for the therapeutic group on 5 dpi. And the livers, spleens, and lungs were collected for viral load analyses via qRT-PCR.

In the prophylactic treatment group, most mice died between 5-6 dpi without mAbs injection and exhibited significantly higher viral RNA copies in serum samples (Figures 3D and 3E). BD70-4003 (ⅠA), BD70-4008 (ⅢA), and BD70-4017 (ⅢA) protected 100% mice from SFTSV infection, while one mouse died on day 7 or day 10 in the groups treated with BD70-4009 (ⅢA), BD70-4013 (ⅠA), and BD70-4022 (ⅢA) (Figure 3D). Viral copy numbers in the serum of mice treated with BD70-4003, BD70-4008, or BD70-4017 were below the detection limit, indicating that these three antibodies exhibited superior efficacy in promoting viral clearance (Figure 3E).

In the therapeutic treatment group, BD70-4003 (ⅠA) and BD70-4017 (ⅢA) protected 100% mice from SFTSV challenge (Figure 3G), and extremely low viral copy numbers were detected in the serum samples collected on both 3 or 5 dpi (Figures 3H and 3I). The remaining four antibodies conferred weaker protective effects, with varying numbers of deaths occurring within their respective treatment groups. We also analyzed tissue samples collected on 5 dpi. Viral copy numbers in spleens, livers, and lungs from the BD70-4003 and BD70-4017 treatment groups were reduced by 3-4 log values compared to the vehicle group (Figures 3J, 3K, and 3L). All these results suggested that administration of either BD70-4003 (ⅠA) or BD70-4017 (ⅢA) exhibited significant therapeutic effects. The *in vivo* experimental results further validated our preliminary *in vitro* antibody screening strategy, demonstrating that antibodies targeting epitopes IA and IIIA hold the greatest potential for development as therapeutic antibodies.

### Structural analyses of Gn-Ab complexes

To gain deeper insights into the molecular mechanisms underlying the distinct neutralizing activities of different antibodies, we selected 3 mAbs (BD70-4003, BD70-4008, and BD70-4017) for structural analysis. Among them, BD70-4003 and BD70-4008 exhibited competitive binding to Gn protein. Consequently, except Gn head/BD70-4003 Fab and Gn head/BD70-4008 Fab, we also designed 2 combinations of antibodies to form complexes with the Gn-head for structural analysis, thereby increasing the complex size to facilitate structural determination. The Gn head/BD70-4003 Fab and Gn head/BD70-4008 Fab were detected by cryo-EM (Table S8), with their structure ultimately resolved at reported global resolution with 2.85 and 2.97 Å, respectively (Figures S5A and S5B). Two complexes, Gn head/BD70-4003 Fab/BD70-4017 Fab and Gn head/BD70-4008 Fab/BD70-4017 Fab, were also resolved with both 2.69 Å resolution (Figures S5C and S5D).

The complex structure of the BD70-4003 Fab bound to the Gn-head revealed a large interaction interface primarily located on Domain I. Key contact residues on Gn included S32, K34, Q68, Q73-R74, S145-K147, S158-G161, S163-S164, and L166-S169 (Figures 4A and 4B). The interaction is stabilized by 11 hydrogen bonds, with the heavy chain (specifically residues D55, E100, D101, D104, and Y105) contributing the majority of the bonding potential (Figure 4C). This epitope overlaps significantly with the identified binding sites for the host receptor C-C motif chemokine receptor 2 (CCR2).^40^ Specifically, the proximity to residue D170, which is a critical determinant of SFTSV-Gn binding to CCR2, indicates that BD70-4003 neutralizes the virus by competitively inhibiting receptor attachment.^41^

**Figure 4.**
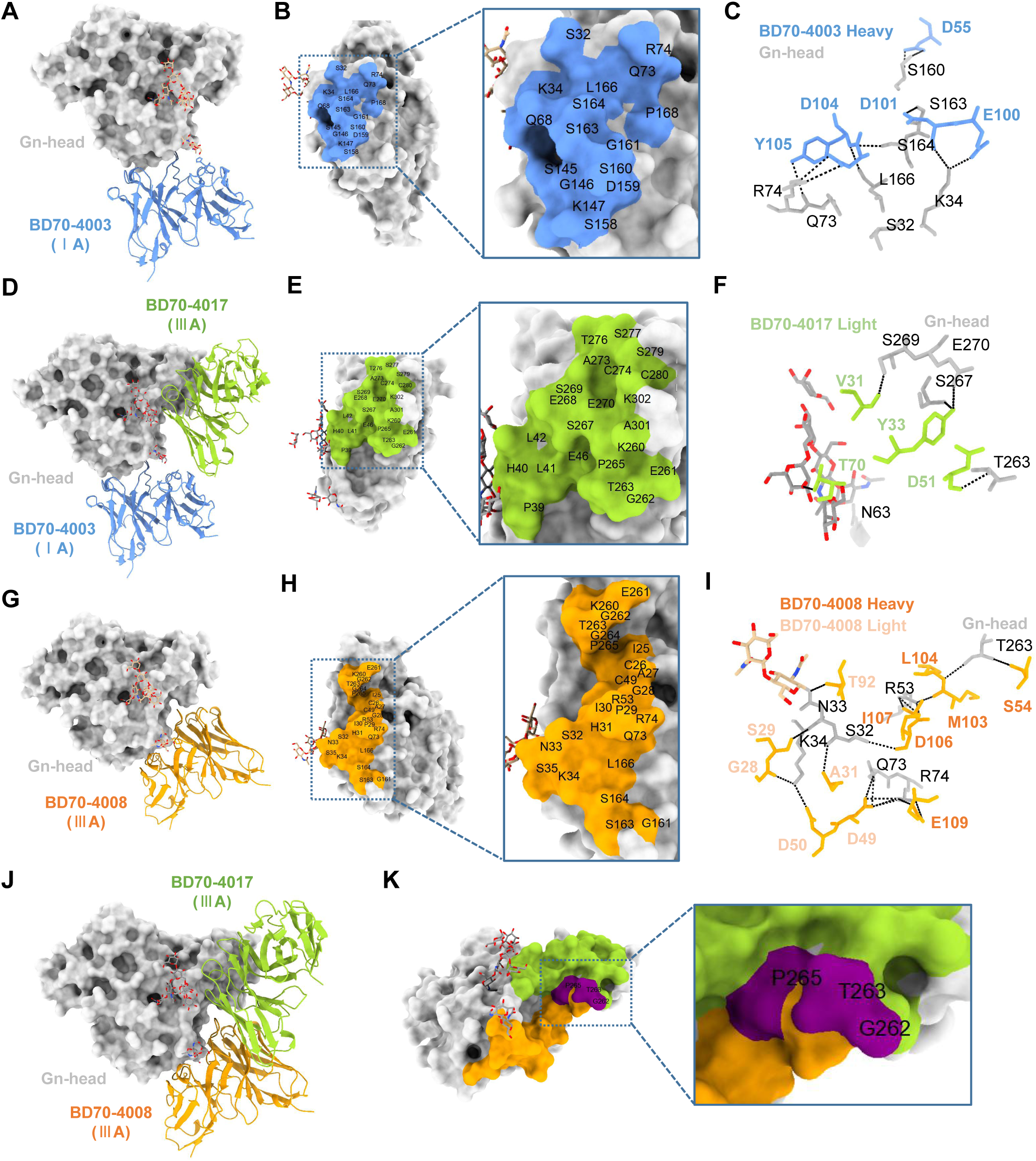
Cryo-EM structure of Gn antibodies bound to HB29-Gn-head. (A, D, G and J) Aligned structures of BD70-4003 (A), BD70-4003/BD70-4017 (D), BD70-4008 (G) and BD70-4008/BD70-4017 (J) on SFTSV Gn-head. BD70-4003, BD70-4008 and BD70-4017 are colored in blue, orange, and green, respectively. Gn-head is colored in gray. (B, E and H) Surface representation of the contact residues on Gn-head interacting with BD70-4003 (B), BD70-4017 (E) and BD70-4008 (H). (C, F and I) Interactions between the heavy-chain and light-chain BD70-4003 (C), BD70-4017 (F) and BD70-4008 (I) and Gn-head. Residues of Gn-head, BD70-4003, BD70-4008 or BD70-4017 are gray, blue, orange, and green, respectively. (K) The overlapping epitopes of BD70-4008 and BD70-4017 are labeled in purple. See also Figures S5-S6 and Table S8.

In contrast, BD70-4017 primarily targets Domain III, engaging residues K260-T263, P265, S267-E270, and A273-C280 (Figures 4D and 4E). A significant feature of the BD70-4017 binding mode is its interaction with the N-linked glycan at position N63 of Domain I (Figure 4F). The light chain residue T70 forms a hydrogen bond with the mannose moiety of this glycan, a mechanism that may enhance the stability of the complex on the authentic virion surface. Given that Domain III acts as a structural cap over the Gc fusion loop, the binding of BD70-4017 likely hinders the exposure of the fusion machinery, thereby preventing the acidification-induced conformational changes required for viral entry.

The structural analysis of BD70-4008 provided a cautionary insight into the interpretation of DMS data. Although DMS clustered BD70-4008 into group IIIA along with BD70-4017, the cryo-EM structure revealed that BD70-4008 actually interacts with a hybrid epitope spanning Domain I and Domain III (Figure 4G-I). The discrepancy arose because BD70-4008 possesses relatively few escape residues (Q50, R74, T263), which made its DMS escape profile computationally similar to the Domain III-focused BD70-4017. This “grouping bias” highlights that while DMS is a powerful tool for rapid classification, it can underrepresent complex, multi-domain epitopes if only a subset of residues provides strong escape signals. Interestingly, although the amino acid residues on the interacting interface of BD70-4008 and BD70-4017 have a slight overlap, involving 3 residues (G262, T263 and P265), the combination of the two antibodies has little hindering effect on the formation of the Gn head/BD70-4008/BD70-4017 complex (Figure 4J-K).

To investigate the neutralization mechanisms of BD70-4003, BD70-4008, and BD70-4017, we superimposed the structures of the BD70-4003/BD70-4017-Gn and BD70-4008/BD70-4017-Gn complexes onto the native SFTSV virion structure. Previous studies have shown that Gn-Gc heterodimers form hexon and penton on the surface of SFTSV virions.^12,42^ Our results showed that BD70-4003/BD70-4017 and BD70-4008/BD70-4017 binding sites were located on the surfaces of the hexons or pentons (Figures 5A and 5B). BD70-4003 or BD70-4008 binds to the top side of the authentic SFTSV virion with an orientation nearly perpendicular to the viral membrane, which may be the key factor contributing to the superior affinity and neutralization effects. Although BD70-4017 binds to a more lateral position, its binding is less affected and experiences minimal steric hindrance. This structural complementarity supports the potential for combining these antibodies into a therapeutic cocktail.

**Figure 5.**
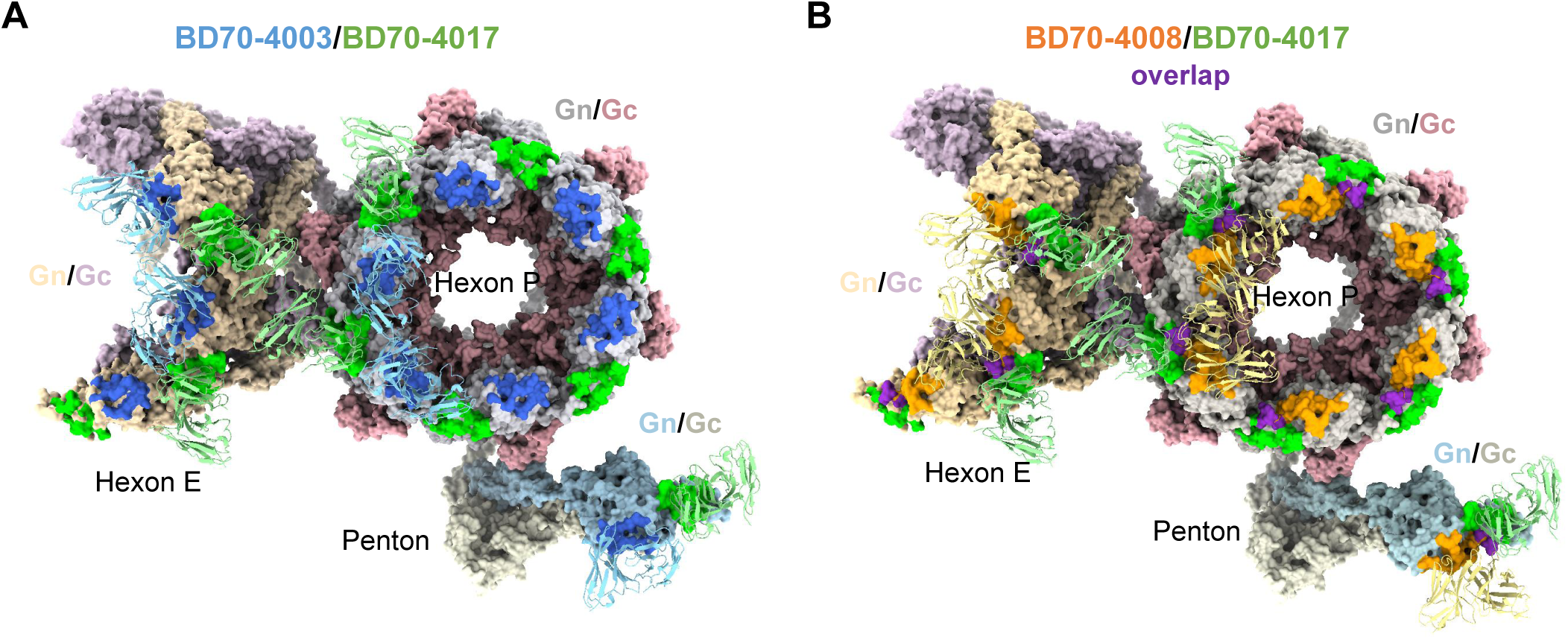
Docking of three antibodies onto the surface of the SFTSV virion. Docking of BD70-4003/BD70-4017 (A) and BD70-4008/BD70-4017 (B) onto the surface of SFTSV virion (PDB: 8I4T). BD70-4003, BD70-4008 and BD70-4017 are colored in blue, orange, and green, respectively. The binding sites of mAbs on SFTSV virion are labeled with colors within the same palette.

## DISCUSSION

Currently, the clinical management of SFTS is centered on comprehensive supportive care, including platelet transfusion, intravenous immunoglobulin, and blood purification.^43–45^ Due to the lack of specific and effective antiviral drugs, severe cases can rapidly progress to multiple organ failure, particularly in older patients, with significantly increased mortality rates.^46^ In this study, we successfully combined pseudovirus screening with authentic virus validation to identify potent mAbs that may serves as SFTSV therapeutics.^32^ Compared to authentic viruses, pseudoviruses offer advantages including lower strain restrictions, higher detection throughput, and operational convenience, making them suitable for preliminary efficacy screening. However, we observed that antibody BD70-4017, which displayed relatively modest performance in the pseudovirus system, demonstrated significantly enhanced neutralizing activity against the authentic virus (Figure 1 and Table S3), highlighting that the VSV-G-based pseudovirus system cannot completely mimic the surface glycoprotein arrangement of authentic SFTSV. Similar limitations have been previously reported in studies on both SFTSV^15^ and hantavirus^47^, and lentiviral-based SFTSV pseudoviruses^48,49^ also appear unable to fully recapitulate authentic viral features. Future research should focus on optimizing existing systems or developing novel platforms that accurately mimic the glycoprotein distribution of the authentic virus, thereby enabling high-throughput antibody testing in conventional laboratory settings.

Mapping antibody epitopes is essential for understanding neutralization mechanisms and guiding rational vaccine and therapeutic design. The implementation of a high-resolution DMS platform for bunyavirus antibody characterization represents a significant technical advancement. Traditional epitope mapping methods, such as competitive ELISA or cross-blocking assays, provide only coarse-grained information about epitope groups. In contrast, DMS provides residue-level specificity, allowing for the immediate identification of potential escape variants and the rational design of antibody cocktails. By comprehensively evaluating the impact of every possible amino acid substitution on antibody binding, researchers can proactively monitor viral antigenic evolution in the field and update therapeutic strategies accordingly. Also, the revision of the SFTSV Gn antigenic landscape into eight specific epitope groups clarifies why previous studies observed varying degrees of protection. For instance, targeting the variable regions of Domain III (IIIB) often leads to strain-specific or weak protection compared to the functional interfaces in Domain III (IIIA). Specifically, the MAb4-5 (IIIB) is thought to neutralize infection by recognizing a transient intermediate state during membrane fusion. In the prefusion state of mature virions, however, this epitope is masked by adjacent Gn/Gc subunits, likely causing steric hindrance that restricts binding and limits neutralizing potency.^12^ In contrast, BD70-4017 (IIIA) is not subject to such steric constraints (Figure 5). These differences may account for the distinct neutralizing activities observed between the two epitope groups.

The identification of the BD70 series coincides with a burgeoning field of SFTSV immunology characterized by the discovery of several other potent human and murine antibodies.^13–17^ Recent studies have introduced several high-performing antibody series, notably the JK series (JK-2, JK-8),^18^ the SD series (SD4, SD12, SD22),^19^ and the ultrapotent ZS1C5^50^. Like BD70-4003, the JK series and SD series primarily target Domain I. SD4 and SD22 exhibit picomolar binding affinities (32–83 pM) and have shown 100% protection in mice when given at 20 mg/kg at 3 dpi, representing a potentially wider therapeutic window than the BD70 series if higher doses are used.

The antibody ZS1C5 represents a distinct class of “interdomain” neutralizers.^50^ Structural analysis has revealed that ZS1C5 bridges Domain I and Domain III of the Gn-head, mediated by an unusually long CDRH3 of over 17 residues. ZS1C5 exhibits subnanomolar potency and has demonstrated robust protection in both murine and non-human primate (NHP) models, the latter being a significant milestone for SFTSV therapeutics. While BD70-4003 and BD70-4017 target more traditional domain-specific epitopes, their high efficacy and distinct binding sites make them excellent partners for cocktails, potentially mirroring the success seen with other emerging interdomain-targeting platforms.

In conclusion, the delineation of the SFTSV Gn antigenic landscape and the functional characterization of human mAbs provides a definitive path forward for both therapy and prevention. The integration of high-throughput mutational screening with atomic-level structural analysis has resolved long-standing questions regarding epitope-function relationships, identifying the Gn-head Domain I and Domain III as the primary targets for neutralizing intervention. The unique structural insights provided by the DMS-cryo-EM synergy not only advance the development of SFTSV-specific drugs but also establish a generalizable model for responding to other emerging tick-borne viruses. These findings, coupled with the demonstration of 100% in vivo protection, provide a robust foundation for the rational design of universal vaccines for SFTSV and its related pathogens.

### Limitations of the study

Our study has established a powerful framework for discovering bunyavirus antibodies by employing DMS and identified promising therapeutic candidates, but some limitations remain. First, the binding sites of some mAbs cannot be efficiently identified by DMS due to weak escape signal or high noise from non-epitope sites, and the DMS-based unsupervised clustering is limited by the size of mAb collection in this study. These epitope mapping results could be complemented by structural analyses. Second, the conclusions derived from pseudovirus-based neutralization assays cannot be always directly translated to inhibition against live-virus or efficacy *in vivo*, highlighting the necessity of further validation. Lastly, the effectiveness of BD70-4003 or BD70-4017 against advanced-stage infections has not been fully demonstrated, since patients typically present for treatment multiple days after symptoms emerge. Further pre-clinical and clinical research should prioritize refining dosing protocols and assessing long-term safety and efficacy.

## RESOURCE AVAILABILITY

### Lead contact

Further information and requests for resources and reagents should be directed to and will be fulfilled by the lead contact, (Y.C.).

### Materials availability

All unique reagents generated in this study are available from the lead contact with a completed Material Transfer Agreement.

### Data and code availability

All data and materials presented in this manuscript are available from the lead contact upon a reasonable request under a completed Material Transfer Agreement. This paper does not report original code. Any additional information required to reanalyze the data reported in this paper is available from the lead contact upon request.

## ACKNOWLEDGMENTS

We thank Yantai Qishan Hospital for the assistance in recruiting SFTSV-infected convalescents and collecting samples. We thank Yuhang Wang from Changping Laboratory for the support of structure analysis. We thank Nanjing GenScript Biotechnology for the technical assistance on the purification of mAbs. This project is financially supported by Changping Laboratory (2025D-04-01).

## AUTHOR CONTRIBUTIONS

Y.C. designed the study. Y.C., Q.W., and J.L. wrote the manuscript with input from all authors. Y.C. and F.S. coordinated the expression and characterization of the neutralizing antibodies. F.J., Q.W., and H.S. performed and analyzed the yeast display screening experiments. Q.W., Y.Y., J.W., L.Y., and Y.W. performed the neutralizing antibody expression and characterization, including pseudovirus neutralization assays and ELISA. Q.W., M.M, and F.J. performed and analyzed the antigen-specific single B cell VDJ sequencing. Y.W. and Q.W. performed the structural analyses. W.L., H.L. and A.H. performed and coordinated the authentic virus neutralization and animal experiments. Y.L. and L.Z. recruited the SFTSV convalescents.

## SUPPLEMENTAL INFORMATION

**Document S1. Figures S1–S6, Tables S1–S2, S4-S5, S7-S8**

**Table S3. Neutralizing activities of SFTSV-Gn or Gc antibodies, related to Figure 1**

**Table S6. Germline and CDR3 sequences of SFTSV-Gn or Gc antibodies, related to Figure 2**

## METHODS

### EXPERIMENTAL MODEL AND STUDY PARTICIPANT DETAILS

#### Cells

For neutralization assays, Huh-7 cells and Vero cells were cultured in DMEM supplemented with 10% FBS, maintained at 37℃ in an incubator supplied with 8% CO2.

#### SFTSV VSV-based pseudovirus

The SFTSV pseudovirus was constructed as previously described using VSV pseudotyped virus (G*ΔG-VSV).^32^ Pseudovirus carrying glycoprotein of SFTSV HB29, HN20, HN13, SPL030A, SD4, G/K/2012, WCH, HBMC16, and AHL were constructed and used, as described previously.^32^

#### Authentic SFTSV virus

SFTSV HBMC16, WCH/97/HN/China/2011, HNXY2017-50, and HNXY2017-66 were stored in the State Key Laboratory of Pathogen and Biosecurity of Academy of Military Medical Sciences. The virus load of the cells was determined using a standard 50% focus reduction neutralization test FRNT_50_ assay.

#### IFNAR-/- mice

Specific-pathogen-free, 6 to 8-week-old IFNAR-/- C57BL6/J mice (18-20g) were raised by the State Key Laboratory of Pathogen and Biosecurity of Academy of Military Medical Sciences as previously described.^18^ Animal experiment was performed in accordance with the National Institutes of Health guidelines under protocols approved by the Institutional Animal Care and Use Committee (IACUC-IME-2021-003).

## METHOD DETAILS

### Plasma donors

Blood samples were collected from 12 convalescent individuals previously infected with SFTSV. All plasma-related experiments were approved by the Ethics Committee of Beijing Ditan Hospital, Capital Medical University (approval no. DTEC-KY2022-022-01). Written informed consent was obtained from all participants in accordance with the Declaration of Helsinki, permitting the collection, storage, and research use of clinical samples, as well as publication of resulting data. Whole blood samples were diluted 1:1 with PBS containing 2% FBS and subjected to Ficoll (Cytiva, 17-1440-03) density gradient centrifugation to separate plasma and PBMCs. Plasma was collected from the upper layer, while cells at the interface were harvested for PBMC isolation. PBMCs were further processed by centrifugation, red blood cell lysis (Invitrogen eBioscience 1× RBC Lysis Buffer, 00-4333-57), and washing steps. Samples not used immediately were were stored in FBS (Gibco) containing 10% DMSO (Sigma) in liquid nitrogen. Cryopreserved PBMCs were thawed in DPBS containing 2% FBS (Stemcell, 07905) prior to use.

### Antigen-specific cell sorting, V(D)J sequencing, and data analysis

PBMCs were processed using the EasySep Human CD19 Positive Selection Kit II (STEMCELL, 17854) to isolate CD19^+^ B cells. Cells were stained with FITC anti-human CD19 (BioLegend, 302206, clone HIB19), Brilliant Violet 421 anti-human CD27 (BioLegend, 302824, clone O323), and PE/Cyanine7 anti-human IgM (BioLegend, 314532, clone MHM-88). For antigen labeling, biotinylated SFTSV HB29-Gn or HB29-Gc protein was conjugated with PE-streptavidin (BioLegend, 405204) and APC-streptavidin (BioLegend, 405207), respectively. Biotinylated ovalbumin (OVA) conjugated with streptavidin served as a negative control. Specifically, HB29-Gn and HB29-Gc were labeled with distinct barcodes: Gn was conjugated with TotalSeq™-C0971 and TotalSeq™-C0972 Streptavidin (BioLegend, 405271 and 405273), while Gc was conjugated with TotalSeq™-C0973 and TotalSeq™-C0974 Streptavidin (BioLegend, 405275 and 405277) to enable sequencing-adapted sorting. OVA was labeled with TotalSeq™-C0975 Streptavidin (BioLegend, 405279). Cells were incubated on ice in the dark for 30 minutes and washed twice. Dead cells were excluded by 7-AAD staining (Invitrogen, 00-6993-50). 7-AAD⁻, CD19⁺, CD27⁺, IgM⁻, OVA⁻, and antigen-positive (Gn⁺ or Gc⁺) B cells were sorted using a MoFlo Astrios EQ Cell Sorter (Beckman Coulter). FACS data were acquired with Summit 6.0 (Beckman Coulter) and analyzed using FlowJo v10.8 (BD Biosciences).

Sorted cells were processed using the 10X Genomics Chromium Next GEM Single Cell V(D)J kit (v1.1, CG000208) according to the manufacturer’s protocol to generate barcoded libraries via GEM formation, reverse transcription, and cDNA amplification. Libraries were sequenced on an Illumina platform.

BCR contigs were assembled and aligned to the reference using Cell Ranger (v6.1.1). Only productive contigs and B cells with paired heavy and light chains were retained after quality filtering. Germline V(D)J gene annotation was performed with IgBlast (v1.17.1), and somatic hypermutations were analyzed using the Change-O toolkit (v1.2.0).

### Monoclonal antibody expression and purification

Human antibody heavy- and light-chain genes were synthesized and cloned into respective pCMV3 vectors (pCMV3-CH, pCMV3-CL, or pCMV3-CK). Following transformation into E. coli DH5α (Tsingke, TSC-C01-96) and overnight incubation at 37°C, positive clones were identified by colony PCR and Sanger sequencing, followed by plasmid preparation (CWBIO, CW2105). For protein expression, Expi293F cells were co-transfected with paired plasmids using polyethylenimine (Yeasen, 40816ES03) in 0.9% NaCl. Transfected cultures were maintained under standard growth conditions, with nutrient supplement (OPM Biosciences, F081918-001) added 24 h post-transfection and replenished every 48 h. Supernatants were harvested after 8 days of culture for antibody purification.

Antibodies were purified from clarified supernatant (3,000 × g, 10 min) using Protein A magnetic beads (GenScript, L00695) and a KingFisher system (Thermo Fisher). Final samples were quantified by NanoDrop (Thermo Fisher, 840-317400) and analyzed for purity by SDS-PAGE (LabLead, P42015).

### ELISA

Purified SFTSV Gn or Gc proteins (1 μg/mL in Solarbio C1055 buffer) were used to coat ELISA plates overnight at 4°C. After washing and blocking, serially diluted antibodies were added and incubated for 30 minutes at room temperature. The plates were then incubated with an HRP-conjugated goat anti-human IgG (Jackson ImmunoResearch, 109-035-003; 0.25 μg/mL) for 30 minutes, followed by tetramethylbenzidine (TMB) substrate (Solarbio, PR1200) for signal development. The reaction was terminated with H₂SO₄, and absorbance at 450 nm was read on a microplate reader (Multiskan Fc, Thermo Scientific).

### Surface Plasmon Resonance

SPR experiments were performed on a Biacore 8K (Cytiva). SFTSV mAbs (human IgG1) were captured by a Sensor Chip Protein A (Cytiva). Various concentrations of SFTSV Gn (His-tag; 1.5625, 6.25, 12.5, 25, and 50 nM) were injected. The response was recorded at room temperature, and the raw data curves were fitted to a 1:1 binding model using Biacore Insight Evaluation Software (Cytiva, v4.0.8).

### Pseudovirus neutralization assay

SFTSV Gn/Gc pseudoviruses were produced using a VSV-based system. Briefly, 293T cells were infected with G*ΔG-VSV (Kerafast) and transfected with a plasmid encoding the SFTSV glycoproteins. The culture supernatant containing the pseudoviruses was collected, clarified by filtration, aliquoted, and stored at -80°C. For the neutralization assay, serially diluted monoclonal antibodies were mixed with pseudoviruses and incubated for 1 hour at 37°C with 5% CO₂. Huh-7 cells were then added to the mixture. After 48 hours, the supernatant was removed, and luminescence was measured following the addition of D-luciferin substrate (PerkinElmer, 6066769) using a microplate reader (PerkinElmer, HH3400). The half-maximal inhibitory concentration (IC₅₀) was determined by fitting the data to a four-parameter logistic model in GraphPad Prism (v9.0.1).

### DMS library construction

DMS libraries were constructed as previously described, with modifications for the SFTSV Gn-head domain (residues 20-340). Using the HB29 Gn-head coding sequence as a template, two independent mutant libraries were generated via three rounds of mutagenesis PCR with synthesized primer pools. Each Gn-head variant was constructed with an N-terminal HA tag and a C-terminal MYC tag, linked to a unique 26-nucleotide barcode. Barcode-variant mapping was performed by PacBio sequencing. The mutant libraries were cloned into the pETcon vector and electroporated into DH10B cells for plasmid amplification. Plasmids were then transformed into the EBY100 strain of Saccharomyces cerevisiae as previously reported.^53^ Transformed yeast were selected on SD-CAA plates and expanded in SD-CAA liquid medium. Yeast libraries were flash-frozen in liquid nitrogen and stored at -80°C.

### High-throughput antibody-escape profiling

Antibody escape mutations were assessed using a FACS-based workflow. Yeast-expressing Gn-head mutants were induced overnight and stained. After incubation with primary Gn mAbs for 30 minutes at 4°C, cells were stained with secondary antibodies: APC anti-HA.11 (BioLegend, 901524), FITC-conjugated chicken anti-c-Myc (icllab, CMYC-45F), and PE-conjugated goat anti-human IgG (Jackson, 109-115-098) for 30 minutes at 4°C. Double-positive (MYC⁺ HA⁺) yeast displaying bound mAbs were gated, and mAb-unbound populations adjacent to the main population were sorted and collected. Plasmid DNA was extracted after 40 hours of culture, barcodes were PCR-amplified, purified with AMPure XP beads (Beckman Coulter, A63882), and subjected to high-throughput sequencing on an Illumina NovaSeq X Plus platform.

### Processing of DMS data

Illumina single-end sequencing reads from Gn-head DMS libraries were processed as described with minor modifications. Raw reads were trimmed to 26 bp and mapped to a barcode-variant dictionary using the dms_variants package (v0.8.9). Variant escape scores were calculated as *F × (n_X,ab_/N_ab_)/(n_X,ref_/N_ref_)*, where *n_X,ab_* and *n_X,ref_* is the number of reads representing variant *X*, and *N_ab_* and *N_ref_* are the total number of valid reads in antibody-selected (ab) and reference (ref) library, respectively. *F* is a normalization factor defined as the 99th percentile of escape fraction ratios. Variants with fewer than six reads in the reference library or those associated with severely impaired Gn-head expression were excluded. Global epistasis models were fitted using dms_variants to infer per-mutation escape scores. Escape scores were averaged across replicates that passed quality control.

Site-level escape scores were computed as the sum of mutation escape scores per Gn-head residue. These scores were used as antibody-specific features to generate a matrix ***A****_N_*_×*M*_, where N is the number of antibodies and M is the number of valid positions. Non-surface residues on Gn-head were removed from the analysis. We calculated the pairwise similarities, defined as the square root of the Jensen-Shannon divergence (scipy, v1.11.2) between the feature vectors of two antibodies. The dissimilarity (1.0 - similarity) matrix were used to build a k-nearest neighbor graph (k = 6) and clustered into epitope groups using Leiden algorithm (leidenalg, v0.10.2) with resolution=2.5. These clusters are then refined by integrating structural mapping, V(D)J germline usage, and neutralization data to define biologically meaningful epitope groups.

### Authentic virus neutralization assay

Neutralizing activity of mAbs against authentic SFTSV was measured using a FRNT. The four representative SFTSV strains used for testing include HBMC16, WCH, HNXY2017-50, and HNXY2017-66.^18^ Briefly, serially diluted mAbs were incubated with SFTSV virions for 1 hour at 37°C. The mixture was added to Vero cells and incubated for 1.5 hours at 37°C. Cells were then cultured for 3.5 days in DMEM containing 1.25% methylcellulose. After fixation with 3.7% paraformaldehyde for 30 minutes, cells were stained with an anti-SFTSV NP primary antibody, followed by an HRP-conjugated anti-mouse secondary antibody (Applygen, C1308). Staining was developed using a DAB substrate (TIANGEN, PA110). Plaque numbers were recorded, and FRNT_50_ values were calculated.

### *In vivo* SFTSV Challenge Experiment

Mouse studies were conducted as previously described.^18^ To assess the prophylactic efficacy, *IFNAR1^−/−^* mice received a single intraperitoneal (IP) injection of 5 mg/kg of each mAb or a human IgG1 isotype control antibody one day before IP inoculation with 20 FFUs of SFTSV HBMC16. Mice were monitored daily for clinical signs, body weight, and mortality for over 12 days. Weight data are expressed as percentages of initial weight. Serum samples were collected at 3 days post-infection (dpi) for viral load analysis. The therapeutic assessment was largely identical to the prophylactic experiment described above, except that administration occurred one day post-infection. Serum was collected at 3 and 5 dpi for viral load quantification. At 5 dpi, four mice per group were euthanized, and spleen, lung, and liver samples were harvested for viral load determination.

### RT-qPCR assay

Total RNA from tissues was extracted using the RNAprep Pure Cell Kit (TIANGEN, DP430). Intracellular viral RNA was quantified by RT-qPCR targeting the SFTSV L segment with primes: SFTSV-L-F: 5′-CTCACTCATGCCCTCAACGA-3′ and SFTSV-L-R: 5′-GATGAACTCACCAGCCCTGC-3′. For serum samples, RNA was extracted with the TIANamp Virus RNA Kit (TIANGEN, DP315-R), and viral load was quantified using primers targeting the SFTSV S segment: SFTSV-S-F: 5′-AGCCTAATTGGATATGTCAAATTGC-3′ and SFTSV-S-R: 5′-CGGGTGAAGTGGCTGAAGG-3′.

### Protein expression and purification for structural analysis

Expression constructs encoding the ectodomains of SFTSV Gn or Gc were transiently transfected into HEK293F cells using polyethylenimine (Polysciences). Culture supernatants were harvested 7 days post-transfection, concentrated, and exchanged into binding buffer (25 mM Tris-HCl, pH 8.0, 200 mM NaCl). Proteins were first purified by Ni-NTA affinity chromatography, followed by size-exclusion chromatography on a Superose 6 Increase column in final buffer (20 mM HEPES, pH 7.2, 150 mM NaCl). The Gn-head domain was expressed and purified similarly. Antibody Fabs were expressed and purified as previously described.^54^

### Cryo-EM sample preparation, data collection and model building

For the preparation of the complexes, the purified HB29-Gn-head was mixed with a 1.5-fold molar excess of fab BD70-4003 and BD70-4017, BD70-4008-fab and BD70-4017, BD70-4003, BD70-4008-fab separately, incubated on ice for 40 min and injected onto a Superdex 200 Increase 5/150 column (Cytiva) equilibrated with buffer 1× PBS. SDS-PAGE analysis confirmed the formation of the HB29-Gn-head-3-17, HB29-Gn-head-8-17, HB29-Gn-head-3, HB29-Gn-head-8 complexes in a stoichiometric ratio respectively.

An aliquot of 4 μL protein sample of complex at a protein concentration of 0.5 mg/mL was loaded onto a glow-discharged 300 mesh grid (Quantifoil Au R1.2/1.3). The grids were blotted with a filter paper at 4 ℃ and 100% humidity,^55^ and flash-cooled in liquid ethane using a Thermo Fisher Vitrobot Mark IV and screened using a 200 KV Talos Aectica.

Cryo-EM datasets were collected on a 300kV Thermo Fisher Titan Krios G4 electron microscope equipped with a Falcon 4 camera and a selectris X energy filter (GIF: a slit width of 10eV). The micrographs were collected at a calibrated magnification of x130,000 using the EPU software (Thermo Fisher Scientific), yielding a pixel size of 0.95 Å at object scale. Movies were recorded at an accumulated electron dose of 60e^-^Å^-2^ s^-1^ on each micrograph that was fractionated into a stack of 40 frames with a defocus range of -1.0 μm to -2.0 μm.

Data processing was performed using cryoSPARC (v4.7.1).^56^ The data underwent several steps including Motion Correction, CTF Estimation, Create Templates, Template Picker, Extract from Micrographs, 2D classification, 2D selection for Ab-initio Reconstruction, and subsequent Homogeneous Refinement. To enhance the density around the HB29-Gn-head-Fab region, local refinement was conducted using UCSF ChimeraX (v1.7)^57^ and cryoSPARC. Structural modeling and refinement were performed using WinCoot (v0.9.4.1)^52^ and Phenix (v1.21.2).^51^ Figures were generated using UCSF ChimeraX (v1.7).^58^

## QUANTIFICATION AND STATISTICAL ANALYSIS

SPR assays were performed in two biological replicates. Binding data were analysed with the Biacore Insight Evaluation Software using a 1:1 binding model. Neutralization assays were performed in at least two biological replicates. The half-maximal inhibitory/neutralization concentrations (IC_50_/FRNT_50_) were derived from curve fitting based on a four-parameter logistic regression model.

