## Supplemental information for "Human monoclonal antibodies that target the SFTSV glycoprotein Gn head from four neutralizing epitope groups"

### Epitope and functional classification of human neutralizing antibodies against SFTSV Gn

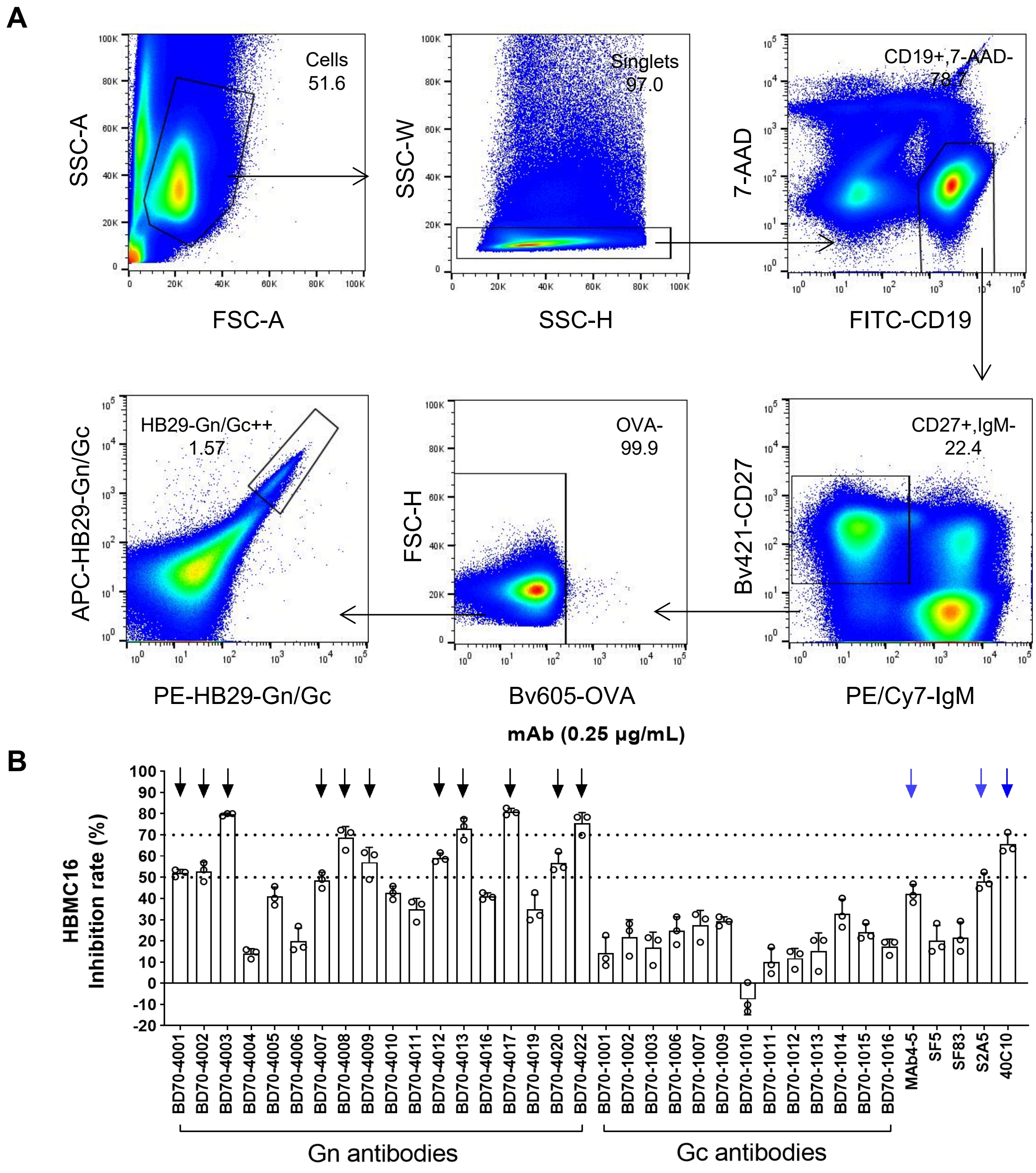


**Figure S1. Isolate memory B cells from SFTSV infected convalescent plasma and screening of mAbs**

(A) FACS strategy to isolate memory B cells. Target of each step and percentage of cells are labeled in each panel.

(B) The inhibition rate of Gn or Gc antibodies against SFTSV HBMC16 authentic viruses at a dose of 0.25μg/ml. Black arrows denote antibodies with superior inhibitory effects, while blue arrows represent control antibodies exhibiting similarly strong inhibition. The inhibition rate were measured in three biological replicates.

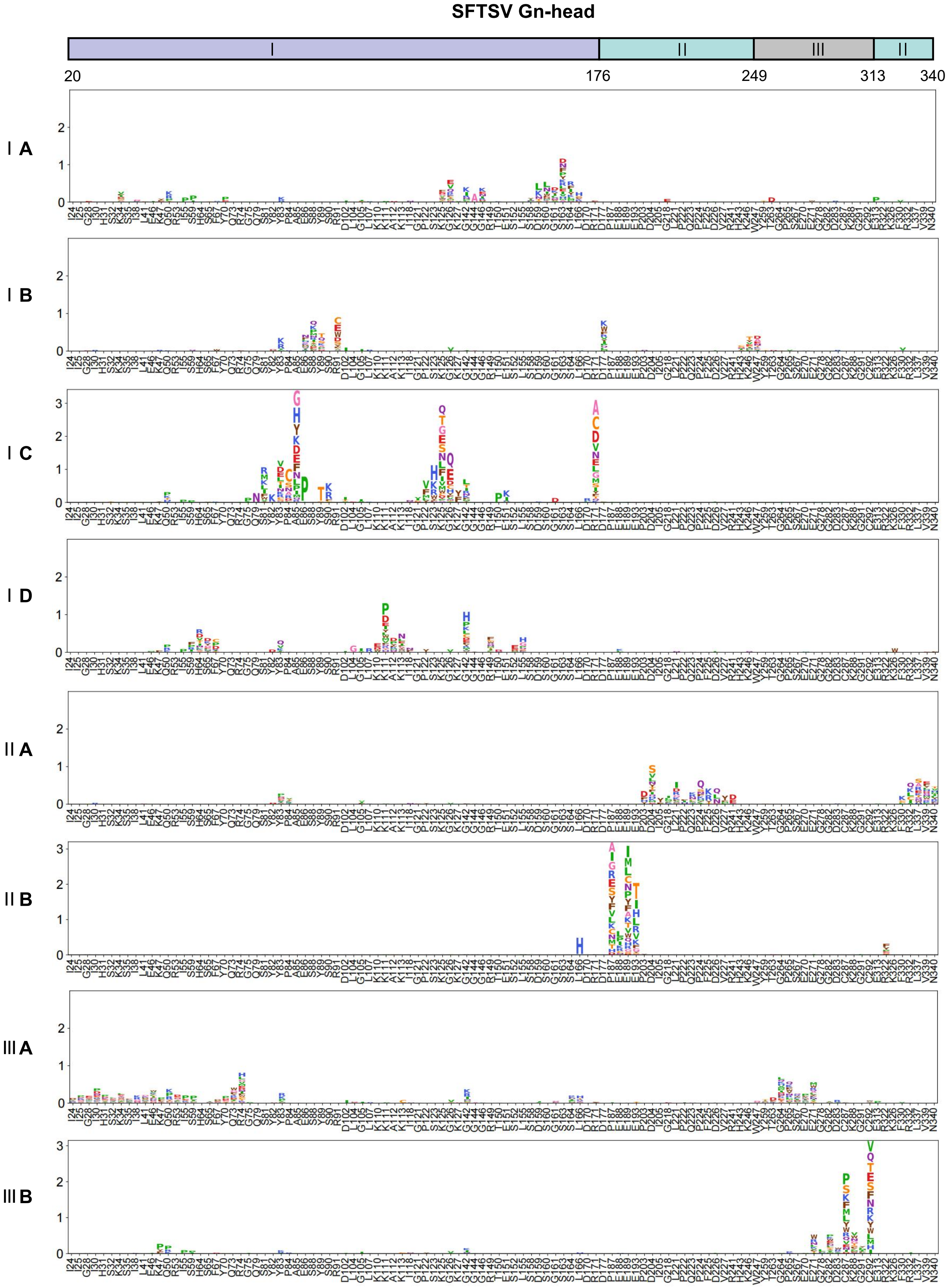


**Figure S2. Average escape scores of antibodies in eight groups**

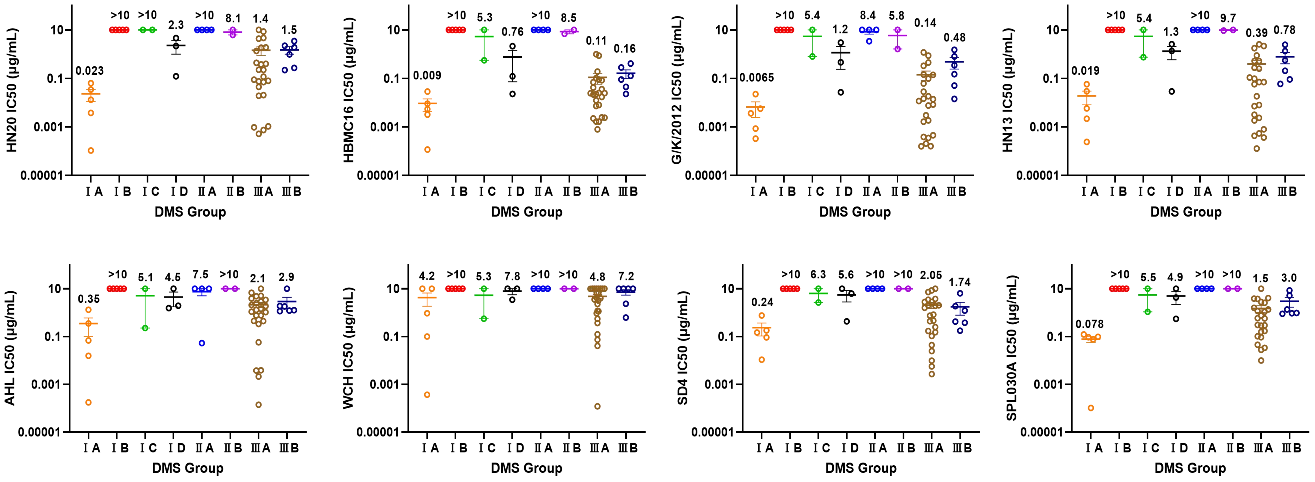


**Figure S3. Statistical analysis of the $IC_{50}$ values against eight SFTSV strains.**

A statistical analysis was performed based on the initial neutralization screening results for eight pseudovirus strains and the DMS-based grouping of each antibody.

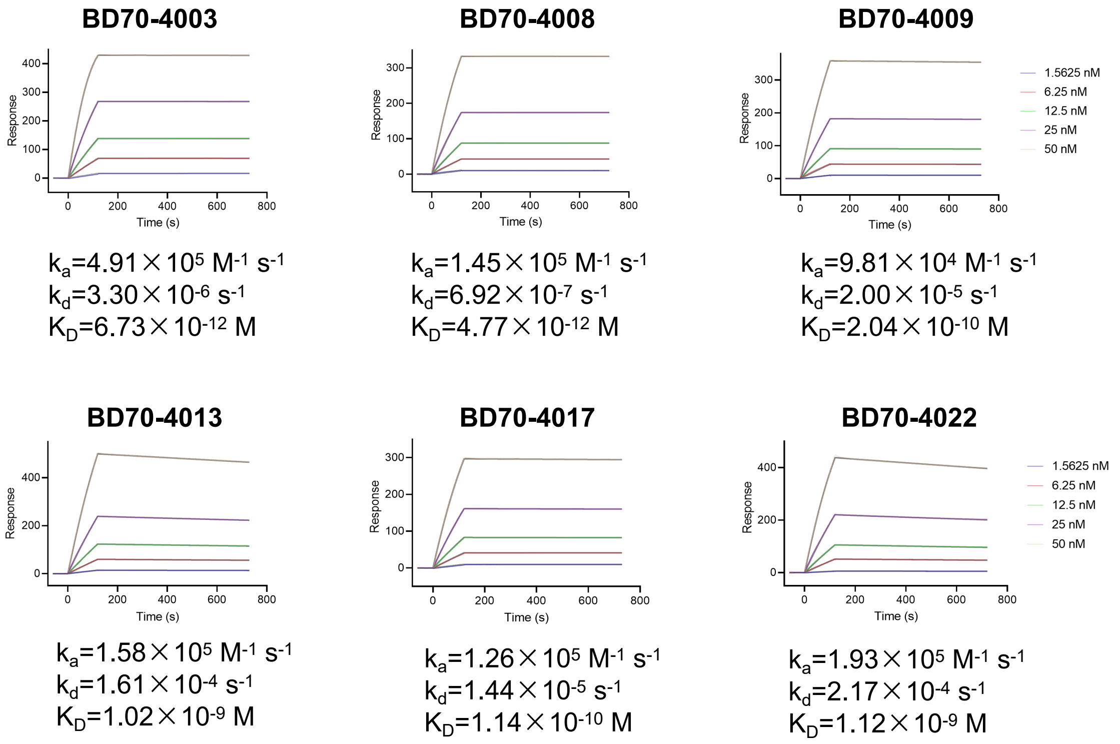


**Figure S4. Antigen affinity of Gn-reactive mAbs**

SPR sensorgrams for the measurement of the binding kinetics of six mAbs to SFTSV Gn protein.

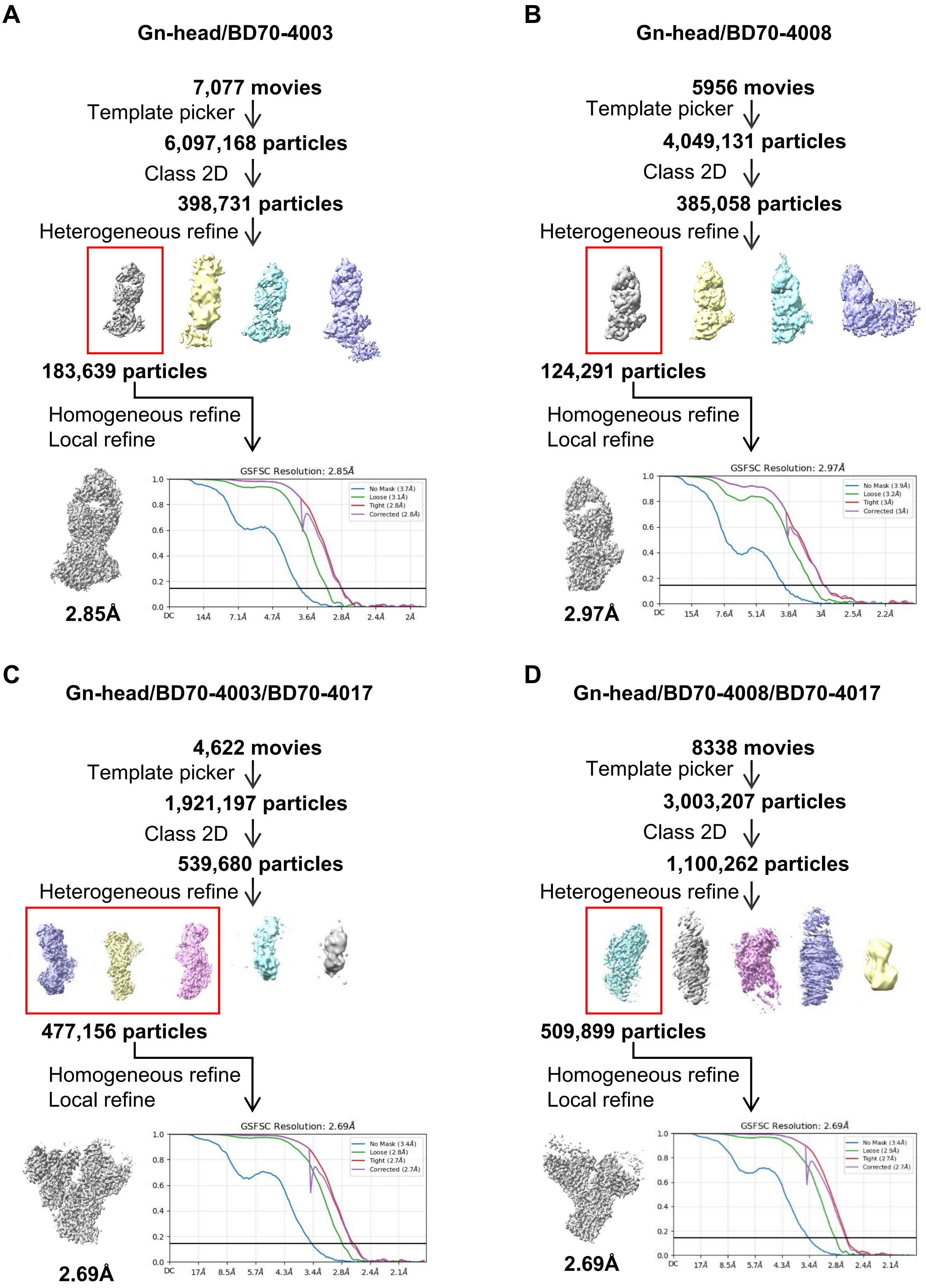


**Figure S5. Workflow for the cryo-EM 3D reconstruction**

Cryo-EM data collection and processing workflow for the reconstruction of the structures of (A) Gn-head/BD70-4003 complex; (B) Gn-head/BD70-4008 complex; (C) Gn-head/BD70-4003/BD70-4017 complex; (D) Gn-head/BD70-4008/BD70-4017 complex.

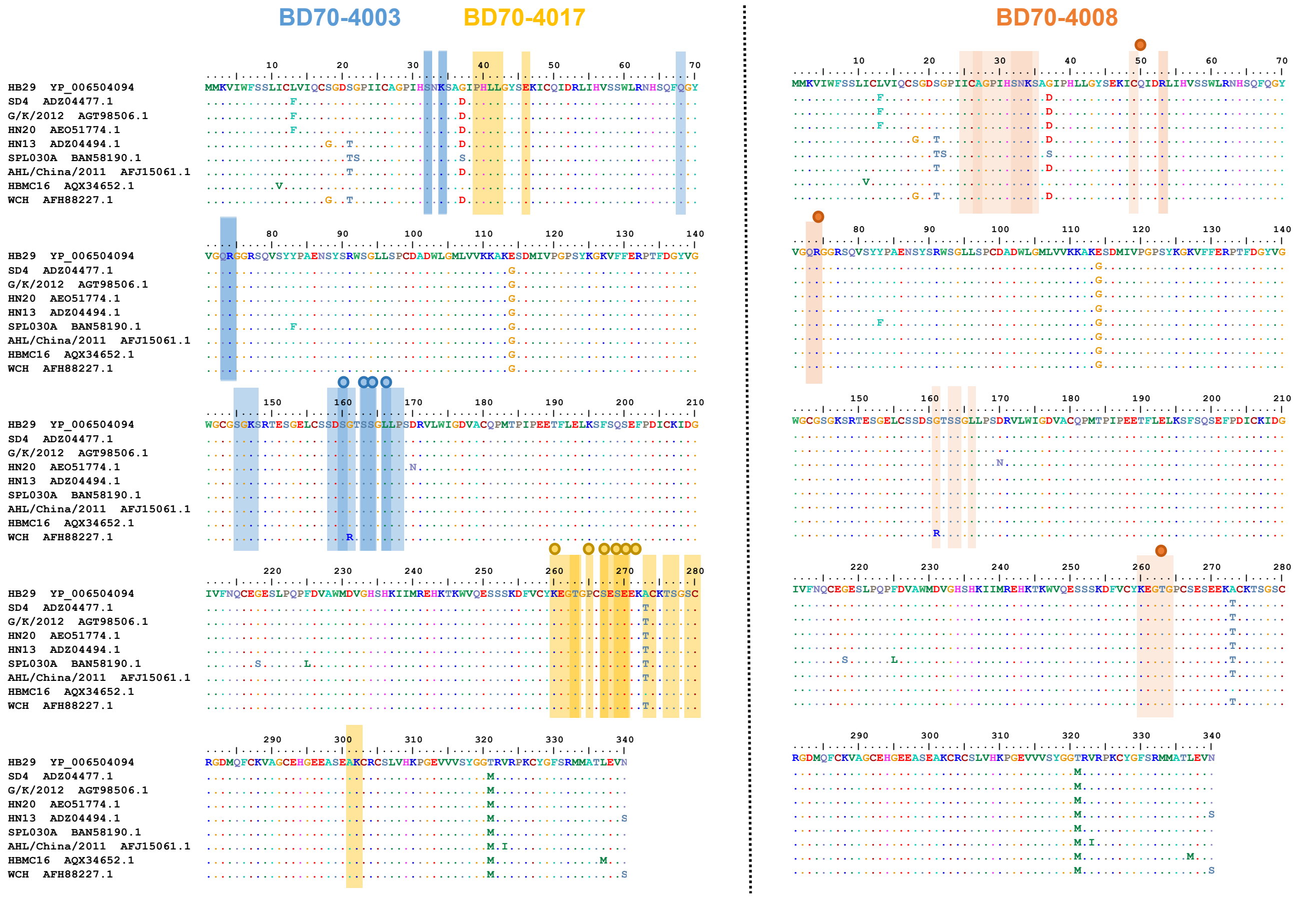


**Figure S6. Sequence alignment of BD70-4003, BD70-4008, and BD70-4017 epitopes**

Different background colors represent distinct antibodies. Light-colored areas indicate amino acids located on the interaction interface, dark-colored areas represent residues involved in binding interactions by cryo-EM. The key amino acid sites identified through DMS are marked with circular dots.

**Supplementary Table 1. Summarized information of SFTSV-infected convalescents**

| Number | Gender | Age | Date of Onset | Date of Admission | Date of Discharge | Date of Collection |
|---|---|---|---|---|---|---|
| SFTS-4 | male | 59 | 2022/8/15 | 2022/8/22 | 2022/9/7 | 2023/11/17 |
| SFTS-5 | female | 54 | 2022/8/16 | 2022/8/23 | 2022/9/7 | 2023/11/17 |
| SFTS-6 | male | 67 | 2022/6/5 | 2022/6/12 | 2022/6/27 | 2023/11/17 |
| SFTS-7 | female | 51 | 2022/7/7 | 2022/7/12 | 2022/7/26 | 2023/11/17 |
| SFTS-8 | female | 69 | 2022/7/16 | 2022/7/21 | 2022/7/31 | 2023/11/17 |
| SFTS-10 | male | 79 | 2022/9/23 | 2022/9/28 | 2022/10/26 | 2023/11/17 |
| SFTS-17 | male | 67 | 2022/7/14 | 2022/7/21 | 2022/9/24 | 2023/11/17 |
| SFTS-18 | female | 62 | 2022/6/25 | 2022/7/1 | 2022/7/20 | 2023/11/17 |
| SFTS-42 | male | 69 | 2022/8/6 | 2022/8/10 | 2022/8/21 | 2023/11/17 |
| SFTS-53 | female | 54 | 2022/6/19 | 2022/6/26 | 2022/7/5 | 2023/11/17 |
| SFTS-58 | female | 54 | 2022/6/25 | 2022/6/29 | 2022/7/8 | 2023/11/17 |
| SFTS-59 | male | 63 | 2022/7/1 | 2022/7/7 | 2022/7/14 | 2023/11/17 |

**Supplementary Table 2. Binding capabilities of SFTSV-Gn or Gc antibodies**

| Antibody ID | Combined Antigen | ELISA OD450 | | | |
|---|---|---|---|---|---|
| | | HB29-Gn 1μg/ml | HB29-Gn 0.1μg/ml | HB29-Gc 1μg/ml | HB29-Gc 0.1μg/ml |
| BD70-4001 | Gn | 3.9982 | 2.01 | - | - |
| BD70-4002 | Gn | 4.209 | 2.5516 | - | - |
| BD70-4003 | Gn | 4.2454 | 3.1048 | - | - |
| BD70-4004 | Gn | 4.4248 | 2.4942 | - | - |
| BD70-4005 | Gn | 3.3408 | 0.769 | - | - |
| BD70-4006 | Gn | 2.4261 | 0.3234 | - | - |
| BD70-4007 | Gn | 4.0894 | 1.8338 | - | - |
| BD70-4008 | Gn | 4.1351 | 3.881 | - | - |
| BD70-4009 | Gn | 3.9192 | 1.3963 | - | - |
| BD70-4010 | Gn | 4.1452 | 2.2829 | - | - |
| BD70-4011 | Gn | 3.3829 | 1.4902 | - | - |
| BD70-4012 | Gn | 3.1742 | 0.6558 | - | - |
| BD70-4013 | Gn | 3.5534 | 1.673 | - | - |
| BD70-4014 | Gn | 4.0679 | 1.4491 | - | - |
| BD70-4015 | Gn | 4.2268 | 1.9839 | - | - |
| BD70-4016 | Gn | 4.1129 | 2.6772 | - | - |
| BD70-4017 | Gn | 4.4014 | 3.6557 | - | - |
| BD70-4018 | Gn | 0.4012 | 0.0365 | - | - |
| BD70-4019 | Gn | 3.383 | 2.0459 | - | - |
| BD70-4020 | Gn | 4.0748 | 1.7407 | - | - |
| BD70-4021 | Gn | 0.4928 | 0.0555 | - | - |
| BD70-4022 | Gn | 3.0257 | 0.8206 | - | - |
| BD70-4023 | Gn | 2.6484 | 0.4138 | - | - |
| BD70-4024 | Gn | 0.151 | 0.0191 | - | - |
| BD70-4025 | Gn | 3.6781 | 1.5736 | - | - |
| BD70-4026 | Gn | 0.2955 | 0.2909 | - | - |
| BD70-4027 | Gn | 4.0611 | 3.9245 | - | - |
| BD70-4028 | Gn | 2.8993 | 0.5814 | - | - |
| BD70-4029 | Gn | 3.9619 | 4.4182 | - | - |
| BD70-4030 | Gn | 4.2833 | 3.8334 | - | - |
| BD70-4031 | Gn | 4.419 | 3.892 | - | - |
| BD70-4032 | Gn | 3.9306 | 3.0193 | - | - |
| BD70-4033 | Gn | 3.0816 | 2.3869 | - | - |
| BD70-4034 | Gn | 4.1666 | 4.8753 | - | - |
| BD70-4035 | Gn | 4.6027 | 4.2137 | - | - |
| BD70-4036 | Gn | 2.3153 | 0.3416 | - | - |
| BD70-4037 | Gn | 0.3249 | 0.0792 | - | - |
| BD70-4038 | Gn | 4.4371 | 4.2241 | - | - |
| BD70-4039 | Gn | 3.9734 | 4.0063 | - | - |

| | | | | | |
|---|---|---|---|---|---|
| BD70-4040 | Gn | 4.3344 | 4.7918 | - | - |
| BD70-4041 | Gn | 3.2681 | 1.5809 | - | - |
| BD70-4042 | Gn | 4.1491 | 3.179 | - | - |
| BD70-4043 | Gn | 4.115 | 3.2518 | - | - |
| BD70-4044 | Gn | 4.3403 | 3.5906 | - | - |
| BD70-4045 | Gn | 3.9281 | 3.6972 | - | - |
| BD70-4046 | Gn | 4.0021 | 3.5577 | - | - |
| BD70-4047 | Gn | 4.3359 | 3.7104 | - | - |
| BD70-4048 | Gn | 4.1934 | 2.8457 | - | - |
| BD70-4049 | Gn | 4.6575 | 5.2975 | - | - |
| BD70-4050 | Gn | 4.2785 | 3.8479 | - | - |
| BD70-4051 | Gn | 4.6539 | 3.9571 | - | - |
| BD70-4052 | Gn | 4.3036 | 4.0624 | - | - |
| BD70-4053 | Gn | 4.219 | 3.5823 | - | - |
| BD70-4054 | Gn | 4.0702 | 2.1892 | - | - |
| 40C10 | Gn | 4.2513 | - | - | - |
| B1G11 | Gn | 4.5718 | - | - | - |
| MAb4-5 | Gn | 4.3986 | - | - | - |
| N1D10 | Gn | 3.9126 | - | - | - |
| S2A5 | Gn | 4.5686 | - | - | - |
| SF1 | Gn | 4.2711 | - | - | - |
| SF5 | Gn | 3.186 | - | - | - |
| BD70-1001 | Gc | - | - | 4.3414 | 3.7413 |
| BD70-1002 | Gc | - | - | 4.3275 | 3.7146 |
| BD70-1003 | Gc | - | - | 4.2212 | 3.8429 |
| BD70-1004 | Gc | - | - | 3.5364 | 0.596 |
| BD70-1005 | Gc | - | - | 0.0529 | 0.0079 |
| BD70-1006 | Gc | - | - | 4.4268 | 3.6198 |
| BD70-1007 | Gc | - | - | 4.3548 | 3.271 |
| BD70-1008 | Gc | - | - | 4.0156 | 0.9029 |
| BD70-1009 | Gc | - | - | 4.5032 | 3.3464 |
| BD70-1010 | Gc | - | - | 4.2999 | 3.4632 |
| BD70-1011 | Gc | - | - | 4.2977 | 2.5328 |
| BD70-1012 | Gc | - | - | 4.0858 | 1.6679 |
| BD70-1013 | Gc | - | - | 4.2928 | 3.3711 |
| BD70-1014 | Gc | - | - | 3.902 | 1.7746 |
| BD70-1015 | Gc | - | - | 4.476 | 3.6973 |
| BD70-1016 | Gc | - | - | 4.1111 | 2.7165 |
| BD70-1017 | Gc | - | - | 0.9325 | 0.0848 |
| BD70-1018 | Gc | - | - | 0.5289 | 0.0556 |
| BD70-1019 | Gc | - | - | 0.1282 | 0.0204 |
| BD70-1020 | Gc | - | - | 0.1216 | 0.0278 |
| BD70-1021 | Gc | - | - | 0.0421 | 0.0148 |
| BD70-1022 | Gc | - | - | 0.0283 | 0.0124 |

| | | | | | |
|---|---|---|---|---|---|
| BD70-1023 | Gc | - | - | 0.0768 | 0.0304 |
| BD70-1024 | Gc | - | - | 0.1401 | 0.0462 |
| BD70-1025 | Gc | - | - | 4.3352 | 4.586 |
| BD70-1026 | Gc | - | - | 5.955 | 5.956 |
| BD70-1027 | Gc | - | - | 2.8109 | 1.2314 |
| BD70-1028 | Gc | - | - | 5.9551 | 5.056 |
| BD70-1029 | Gc | - | - | 4.3714 | 4.3407 |
| BD70-1030 | Gc | - | - | 5.243 | 4.7725 |
| BD70-1031 | Gc | - | - | 4.5399 | 4.2513 |
| BD70-1032 | Gc | - | - | 5.0148 | 4.6131 |
| BD70-1033 | Gc | - | - | 4.3956 | 4.8031 |
| BD70-1034 | Gc | - | - | 4.3985 | 4.0775 |
| BD70-1035 | Gc | - | - | 4.1567 | 3.9116 |
| BD70-1036 | Gc | - | - | 4.4861 | 4.2467 |
| BD70-1037 | Gc | - | - | 4.1203 | 2.8764 |
| BD70-1038 | Gc | - | - | 4.3874 | 4.5896 |
| BD70-1039 | Gc | - | - | 4.4254 | 4.5567 |
| BD70-1040 | Gc | - | - | 4.2205 | 4.0819 |
| BD70-1041 | Gc | - | - | 4.5117 | 4.8136 |
| BD70-1042 | Gc | - | - | 4.0218 | 1.8426 |
| BD70-1043 | Gc | - | - | 4.3429 | 4.3659 |
| BD70-1044 | Gc | - | - | 3.9203 | 4.0622 |

### Supplementary Table 3. Neutralizing activities of SFTSV-Gn or Gc antibodies

| Antibody ID | Epitope Group | Pseudovirus $IC_{50}$ (μg/mL) | | | | | | | | | Inhibition rate (%) against authentic virus at 0.25 μg/mL | | | | Authentic virus $FRNT_{50}$ (μg/mL) | | | |
|---|---|---|---|---|---|---|---|---|---|---|---|---|---|---|---|---|---|---|
| | | HB29 | SD4 | G/K/2012 | HN20 | HN13 | SPL030A | AHL | HBMC16 | WCH | HBMC16 | HNXY 2017-50 | WCH | HNXY 2017-66 | HBMC16 | HNXY 2017-50 | WCH | HNXY 2017-66 |
| BD70-4001 | ID | 0.3201 | >10 | 0.4635 | 2.155 | 1.351 | 4.291 | 1.538 | 0.0227 | >10 | 52 | 38 | 68 | 55 | - | - | - | - |
| BD70-4002 | IIIA | 0.0006042 | 0.2469 | 0.0139 | 0.07887 | 0.003154 | 0.7488 | 0.00376 | 0.007986 | >10 | 52 | 86 | 82 | 73 | - | - | - | - |
| BD70-4003 | IA | 0.0001013 | 0.01072 | 0.0003249 | 0.000105 | 0.0002404 | 0.1214 | 0.0001762 | 0.0001159 | 0.0003676 | 80 | 88 | 88 | 84 | 0.062 | 0.065 | 0.087 | 0.070 |
| BD70-4004 | IIA | >10 | >10 | >10 | >10 | >10 | >10 | 0.05293 | >10 | >10 | - | - | - | - | - | - | - | - |
| BD70-4005 | IIIA | 0.01422 | 8.539 | 0.01844 | 8.523 | 0.1083 | 3.833 | 0.0001392 | 0.02157 | >10 | - | - | - | - | - | - | - | - |
| BD70-4006 | IIIA | 0.2083 | 2.792 | 0.3276 | 4.349 | 0.5207 | 2.567 | 0.05729 | 0.1031 | >10 | - | - | - | - | - | - | - | - |
| BD70-4007 | IIIA | 0.001129 | 0.0437 | 0.008364 | 0.223 | 0.01839 | 0.2837 | 0.003855 | 0.002441 | 0.3657 | 48 | 75 | 72 | 47 | - | - | - | - |
| BD70-4008 | IIIA | 0.0009122 | 0.005597 | 0.0004493 | 0.04444 | 0.0001278 | 0.09376 | 0.00212 | 0.0007943 | 0.5524 | 68 | 83 | 84 | 79 | 0.149 | 0.137 | 0.111 | 0.109 |
| BD70-4009 | IIIA | 0.000385 | 0.009771 | 0.001855 | 0.01907 | 0.007186 | 0.1556 | 3.427 | 0.001658 | 0.07358 | 57 | 85 | 84 | 60 | 0.078 | 0.058 | 0.070 | 0.138 |
| BD70-4010 | IIIA | 0.001144 | 3.439 | 0.09689 | 0.1172 | 0.2723 | 1.346 | 0.4447 | 0.03604 | >10 | - | - | - | - | - | - | - | - |
| BD70-4011 | IC | 0.3266 | 2.66 | 0.8046 | >10 | 0.7489 | 1.07 | 0.2259 | 0.5576 | >10 | - | - | - | - | - | - | - | - |
| BD70-4012 | IA | 0.02317 | 0.754 | 0.0225 | 0.06313 | 0.05719 | 0.09279 | 0.3526 | 0.02848 | >10 | 59 | 78 | 83 | 62 | - | - | - | - |
| BD70-4013 | IA | 0.0003531 | 0.09182 | 0.005605 | 0.03451 | 0.0294 | 0.0001022 | 0.01568 | 0.003193 | 0.1 | 74 | 85 | 90 | 87 | 0.137 | 0.108 | 0.031 | 0.126 |
| BD70-4014 | IIIA | 0.007469 | 2.824 | 0.1403 | 4.624 | 0.8817 | 0.5713 | 1.446 | 0.02427 | 6.832 | - | - | - | - | - | - | - | - |
| BD70-4015 | IB | >10 | >10 | >10 | >10 | >10 | >10 | >10 | >10 | >10 | - | - | - | - | - | - | - | - |
| BD70-4016 | IB | >10 | >10 | >10 | >10 | >10 | >10 | >10 | >10 | >10 | - | - | - | - | - | - | - | - |
| BD70-4017 | IIIA | 4.893 | >10 | 0.1438 | 1.354 | 0.5231 | 1.592 | 6.769 | 0.08962 | 0.04043 | 81 | 80 | 73 | 72 | 0.126 | 0.140 | 0.059 | 0.155 |
| BD70-4019 | IIIA | 0.4599 | 3.748 | 1.178 | 1.88 | 1.768 | 4.051 | 10 | 0.9883 | 6.983 | - | - | - | - | - | - | - | - |

| | | | | | | | | | | | | | | | | | | |
|---|---|---|---|---|---|---|---|---|---|---|---|---|---|---|---|---|---|---|
| BD70-4020 | IIIA | 0.0007605 | 0.1711 | 0.0001593 | 0.001053 | 0.0007945 | 0.04494 | 0.3361 | 0.002375 | 0.00012 | 55 | 75 | 74 | 50 | - | - | - | - |
| BD70-4022 | IIIA | 0.001311 | 0.8177 | 0.003 | 0.07504 | 0.07013 | 0.02801 | 2.087 | 0.02278 | 0.9406 | 78 | 88 | 78 | 77 | 0.111 | 0.231 | 0.234 | 0.143 |
| BD70-4023 | - | 5.82 | >10 | >10 | >10 | >10 | >10 | >10 | >10 | >10 | - | - | - | - | - | - | - | - |
| BD70-4025 | - | >10 | >10 | >10 | >10 | >10 | >10 | 9.208 | >10 | >10 | - | - | - | - | - | - | - | - |
| BD70-4027 | IB | >10 | >10 | >10 | >10 | >10 | >10 | >10 | >10 | >10 | - | - | - | - | - | - | - | - |
| BD70-4028 | IIIA | 0.2124 | 1.97 | 0.8687 | >10 | 2.548 | >10 | 5.066 | 0.8648 | >10 | - | - | - | - | - | - | - | - |
| BD70-4029 | IB | >10 | >10 | >10 | >10 | >10 | >10 | >10 | >10 | >10 | - | - | - | - | - | - | - | - |
| BD70-4030 | IIIA | 0.0002829 | 1.584 | 0.01463 | 0.2259 | 0.09725 | 1.09 | 1.88 | 0.06923 | 1.184 | - | - | - | - | - | - | - | - |
| BD70-4031 | IIA | >10 | >10 | >10 | >10 | >10 | >10 | >10 | >10 | >10 | - | - | - | - | - | - | - | - |
| BD70-4032 | IB | >10 | >10 | >10 | >10 | >10 | >10 | >10 | >10 | >10 | - | - | - | - | - | - | - | - |
| BD70-4033 | IIIA | 0.01875 | 0.4395 | 0.06124 | 0.3749 | 0.07028 | 0.3057 | 1.045 | 0.08373 | 9.842 | - | - | - | - | - | - | - | - |
| BD70-4034 | IA | 0.0004223 | 0.1481 | 0.000847 | 0.003772 | 0.002161 | 0.09721 | 0.06858 | 0.005382 | 0.9385 | - | - | - | - | - | - | - | - |
| BD70-4035 | IIA | >10 | >10 | >10 | >10 | >10 | >10 | >10 | >10 | >10 | - | - | - | - | - | - | - | - |
| BD70-4036 | - | >10 | >10 | >10 | >10 | >10 | >10 | 1.204 | >10 | >10 | - | - | - | - | - | - | - | - |
| BD70-4038 | IIIA | 0.0001698 | 0.02455 | 0.001637 | 0.4362 | 0.002341 | 0.03342 | 2.476 | 0.01719 | 4.17 | - | - | - | - | - | - | - | - |
| BD70-4039 | IIIA | 0.0006396 | 0.1046 | 0.007543 | 0.08815 | 0.007971 | 0.1716 | 0.7686 | 0.001642 | 2.291 | - | - | - | - | - | - | - | - |
| BD70-4040 | IIIA | 0.000327 | 0.002678 | 0.0001589 | 0.0007361 | 0.0003635 | 0.009863 | 0.7549 | 0.007289 | 3.175 | - | - | - | - | - | - | - | - |
| BD70-4041 | IIIA | 0.04024 | 2.01 | 0.446 | 1.845 | 2.159 | 3.739 | 4.841 | 0.1903 | >10 | - | - | - | - | - | - | - | - |
| BD70-4042 | IIIA | 0.0004873 | 1.918 | 0.0003827 | 0.0005125 | 0.0004274 | 0.2975 | 3.896 | 0.02497 | 3.535 | - | - | - | - | - | - | - | - |
| BD70-4043 | IIIA | 0.0001311 | 0.152 | 0.0003483 | 0.0009528 | 0.0005399 | 0.1023 | 1.09 | 0.01238 | 0.1239 | - | - | - | - | - | - | - | - |
| BD70-4044 | IIIA | 0.001037 | 0.4605 | 0.01161 | 0.07038 | 0.05737 | 1.165 | 1.297 | 0.04149 | 4.01 | - | - | - | - | - | - | - | - |
| BD70-4045 | IIIA | 0.0002167 | 0.7472 | 0.0002035 | 0.0204 | 0.001845 | 0.5901 | 0.4485 | 0.002224 | 1.152 | - | - | - | - | - | - | - | - |
| BD70-4046 | IA | 0.0008115 | 0.1801 | 0.003616 | 0.01311 | 0.005812 | 0.07669 | 1.308 | 0.009044 | >10 | - | - | - | - | - | - | - | - |
| BD70-4047 | IIIB | 0.0006097 | 0.4176 | 0.05042 | 0.2191 | 0.09081 | 0.9444 | 1.815 | 0.1265 | 2.356 | - | - | - | - | - | - | - | - |
| BD70-4048 | IIIA | 0.002988 | 7.263 | 0.02827 | 0.3957 | 0.2528 | 2.807 | 2.899 | 0.02459 | >10 | - | - | - | - | - | - | - | - |
| BD70-4049 | IIIB | 0.04375 | 1.22 | 0.8036 | 2.22 | 1.337 | 1.353 | 1.886 | 0.08946 | >10 | - | - | - | - | - | - | - | - |

| | | | | | | | | | | | | | | | | | | |
|---|---|---|---|---|---|---|---|---|---|---|---|---|---|---|---|---|---|---|
| BD70-4050 | IIIB | 0.02536 | 0.3754 | 0.1483 | 0.2712 | 0.2053 | 0.893 | 1.266 | 0.04429 | 0.6172 | - | - | - | - | - | - | - | - |
| BD70-4051 | IIB | >10 | >10 | >10 | >10 | >10 | >10 | >10 | >10 | >10 | - | - | - | - | - | - | - | - |
| BD70-4052 | IIIB | 0.009106 | 1.872 | 0.3456 | 0.9785 | 0.5567 | 5.172 | 1.149 | 0.2802 | >10 | - | - | - | - | - | - | - | - |
| BD70-4053 | - | 0.003816 | 2.066 | 0.1018 | 0.6109 | 0.243 | 1.573 | >10 | 0.04208 | 1.414 | - | - | - | - | - | - | - | - |
| BD70-4054 | - | >10 | >10 | 9.152 | >10 | 8.732 | >10 | 1.988 | >10 | >10 | - | - | - | - | - | - | - | - |
| 40C10 | ID | 0.008162 | 0.4326 | 0.02672 | 0.1216 | 0.02907 | 0.5477 | 1.939 | 0.1224 | 3.464 | 65 | 62 | 83 | 58 | - | - | - | - |
| B1G11 | IIB | 2.797 | 9.99 | 1.63 | 6.189 | 9.402 | >10 | >10 | 6.9 | >10 | - | - | - | - | - | - | - | - |
| MAb4-5 | IIIB | 0.2548 | 6.376 | 1.508 | 3.461 | 2.437 | 8.56 | >10 | 0.4063 | >10 | 42 | 73 | 82 | 60 | - | - | - | - |
| N1D10 | IIA | >10 | >10 | 3.434 | >10 | >10 | >10 | >10 | >10 | >10 | - | - | - | - | - | - | - | - |
| S2A5 | ID | 2.957 | 6.237 | 2.964 | 4.592 | 2.58 | >10 | >10 | 2.122 | >10 | 48 | 21 | 75 | 49 | - | - | - | - |
| SF1 | IIIB | 0.1012 | 0.1755 | 0.01416 | 1.88 | 0.05911 | 0.9679 | 1.205 | 0.02266 | >10 | - | - | - | - | - | - | - | - |
| SF5 | IC | >10 | >10 | >10 | >10 | >10 | >10 | >10 | >10 | >10 | - | - | - | - | - | - | - | - |
| BD70-1001 | - | 3.984 | >10 | >10 | >10 | >10 | >10 | 3.459 | 0.5871 | >10 | - | - | - | - | - | - | - | - |
| BD70-1002 | - | 1.214 | >10 | 7.401 | >10 | >10 | 3.566 | 2.644 | 0.2796 | >10 | - | - | - | - | - | - | - | - |
| BD70-1003 | - | 2.019 | 8.825 | >10 | >10 | >10 | 7.653 | >10 | 0.4965 | >10 | - | - | - | - | - | - | - | - |
| BD70-1004 | - | 7.444 | >10 | 7.066 | >10 | >10 | >10 | >10 | >10 | >10 | - | - | - | - | - | - | - | - |
| BD70-1006 | - | 0.07777 | 0.5713 | 0.714 | 1.379 | 0.8221 | 1.43 | 0.2572 | 0.2317 | 0.3814 | - | - | - | - | - | - | - | - |
| BD70-1007 | - | 3.399 | >10 | >10 | >10 | >10 | >10 | 0.7954 | 0.5881 | 3.205 | - | - | - | - | - | - | - | - |
| BD70-1008 | - | 0.613 | 6.92 | >10 | >10 | >10 | >10 | >10 | >10 | >10 | - | - | - | - | - | - | - | - |
| BD70-1009 | - | 0.2384 | >10 | 6.575 | 3.716 | 3.413 | >10 | >10 | >10 | >10 | - | - | - | - | - | - | - | - |
| BD70-1010 | - | 5.472 | >10 | >10 | >10 | >10 | >10 | 0.8052 | 0.9397 | >10 | - | - | - | - | - | - | - | - |
| BD70-1011 | - | 0.06549 | 0.9009 | 0.8475 | 0.7359 | 0.5571 | 0.6503 | 0.9232 | 0.1423 | 0.4045 | - | - | - | - | - | - | - | - |
| BD70-1012 | - | 0.01431 | 0.1617 | 0.1894 | 0.1985 | 0.1128 | 0.2373 | 0.6824 | 0.04432 | 0.147 | - | - | - | - | - | - | - | - |
| BD70-1013 | - | 0.03296 | 0.3259 | 0.4218 | 0.3443 | 0.3024 | 0.3003 | 0.8866 | 0.1325 | 0.2662 | - | - | - | - | - | - | - | - |
| BD70-1014 | - | >10 | >10 | >10 | >10 | >10 | >10 | >10 | >10 | >10 | - | - | - | - | - | - | - | - |
| BD70-1015 | - | 0.02008 | 0.1764 | 0.2155 | 0.2527 | 0.218 | 0.8002 | 0.3795 | 0.2186 | 0.1321 | - | - | - | - | - | - | - | - |

| | | | | | | | | | | | | | | | | | | |
|---|---|---|---|---|---|---|---|---|---|---|---|---|---|---|---|---|---|---|
| BD70-1016 | - | 0.02766 | 0.3195 | 0.1394 | 0.1727 | 0.1718 | 0.5172 | 0.06199 | 0.0432 | 0.136 | - | - | - | - | - | - | - | - |
| BD70-1025 | - | 1.335 | >10 | >10 | >10 | >10 | >10 | >10 | >10 | >10 | - | - | - | - | - | - | - | - |
| BD70-1026 | - | 0.1421 | 1.192 | 0.5183 | 0.6238 | 0.7641 | 1.811 | 8.159 | 0.9384 | 0.9334 | - | - | - | - | - | - | - | - |
| BD70-1027 | - | 7.56 | >10 | >10 | >10 | 9.102 | >10 | 6.266 | >10 | >10 | - | - | - | - | - | - | - | - |
| BD70-1028 | - | 0.2758 | >10 | 2.11 | 4.828 | 2.759 | >10 | >10 | >10 | 5.658 | - | - | - | - | - | - | - | - |
| BD70-1029 | - | 1.65 | >10 | >10 | 6.689 | >10 | >10 | >10 | 1.771 | >10 | - | - | - | - | - | - | - | - |
| BD70-1030 | - | 1.612 | >10 | 7.509 | 4.233 | 7.777 | >10 | 9.721 | 1.529 | 4.807 | - | - | - | - | - | - | - | - |
| BD70-1031 | - | 0.135 | >10 | 2.257 | 2.015 | 2.05 | >10 | 2.233 | >10 | 2.269 | - | - | - | - | - | - | - | - |
| BD70-1032 | - | 0.01935 | 0.8931 | 0.2698 | 0.2325 | 0.2626 | 0.5997 | 1.071 | 0.1621 | 0.1891 | - | - | - | - | - | - | - | - |
| BD70-1033 | - | 0.01787 | 0.3201 | 0.5035 | 0.4916 | 0.4845 | 1.243 | 0.3581 | 0.1324 | 0.3036 | - | - | - | - | - | - | - | - |
| BD70-1034 | - | 0.5594 | >10 | 3.757 | 4.61 | 5.836 | 7.969 | 3.516 | 1.688 | 1.492 | - | - | - | - | - | - | - | - |
| BD70-1035 | - | 0.4702 | 6.322 | 5.05 | 3.261 | 4.214 | 5.604 | 1.257 | 1.982 | 2.757 | - | - | - | - | - | - | - | - |
| BD70-1036 | - | 0.002436 | 0.3324 | 0.05063 | 0.0471 | 0.09571 | 0.1262 | 3.203 | 0.0764 | 0.2636 | - | - | - | - | - | - | - | - |
| BD70-1037 | - | >10 | >10 | >10 | >10 | >10 | >10 | 1.342 | 9.861 | >10 | - | - | - | - | - | - | - | - |
| BD70-1038 | - | 0.07319 | 1.366 | 0.5503 | 0.5361 | 0.6344 | 0.9422 | 5.855 | 0.5784 | 0.9117 | - | - | - | - | - | - | - | - |
| BD70-1039 | - | 0.2848 | 5.329 | 1.632 | 1.393 | 1.547 | 4.489 | >10 | 1 | 1.941 | - | - | - | - | - | - | - | - |
| BD70-1040 | - | 0.268 | 3.415 | 1.378 | 1.045 | 1.171 | 3.297 | 0.6845 | 1.978 | 1.544 | - | - | - | - | - | - | - | - |
| BD70-1041 | - | 0.5885 | 8.868 | 4.903 | 3.468 | 4.838 | 4.433 | 1.568 | 1.104 | 2.586 | - | - | - | - | - | - | - | - |
| BD70-1042 | - | 2.035 | >10 | >10 | >10 | >10 | >10 | >10 | >10 | >10 | - | - | - | - | - | - | - | - |
| BD70-1043 | - | 0.02637 | 5.483 | 0.3119 | 0.5149 | 0.437 | 3.925 | 9.768 | 2.247 | 1.382 | - | - | - | - | - | - | - | - |
| BD70-1044 | - | 0.0466 | 1.323 | 0.3564 | 0.1979 | 0.5493 | 1.776 | 2.845 | 0.2384 | 0.4537 | - | - | - | - | - | - | - | - |

**Supplementary Table 4. Source data of the heatmap in Figure 1C**

| Lg IC50 (μg/ml) | HB29 | HN20 | HBMC16 | G/K/2012 | HN13 | AHL | WCH | SD4 | SPL030A |
|---|---|---|---|---|---|---|---|---|---|
| BD70-4003 | -3.994 | -3.979 | -3.936 | -3.488 | -3.619 | -3.754 | -3.435 | -1.970 | -0.916 |
| BD70-4020 | -3.119 | -2.978 | -2.624 | -3.798 | -3.100 | -0.474 | -3.921 | -0.767 | -1.347 |
| BD70-4013 | -3.452 | -1.462 | -2.496 | -2.251 | -1.532 | -1.805 | -1.000 | -1.037 | -3.991 |
| BD70-4008 | -3.040 | -1.352 | -3.100 | -3.347 | -3.893 | -2.674 | -0.258 | -2.252 | -1.028 |
| BD70-4002 | -3.219 | -1.103 | -2.098 | -1.857 | -2.501 | -2.425 | >1 | -0.607 | -0.126 |
| BD70-4007 | -2.947 | -0.652 | -2.612 | -2.078 | -1.735 | -2.414 | -0.437 | -1.360 | -0.547 |
| BD70-4040 | -3.485 | -3.133 | -2.137 | -3.799 | -3.439 | -0.122 | 0.502 | -2.572 | -2.006 |
| BD70-4042 | -3.312 | -3.290 | -1.603 | -3.417 | -3.369 | 0.591 | 0.548 | 0.283 | -0.527 |
| BD70-4043 | -3.882 | -3.021 | -1.907 | -3.458 | -3.268 | 0.037 | -0.907 | -0.818 | -0.990 |
| BD70-4034 | -3.374 | -2.423 | -2.269 | -3.072 | -2.665 | -1.164 | -0.028 | -0.829 | -1.012 |
| BD70-4045 | -3.664 | -1.690 | -2.653 | -3.691 | -2.734 | -0.348 | 0.061 | -0.127 | -0.229 |
| BD70-4009 | -3.415 | -1.720 | -2.780 | -2.732 | -2.144 | 0.535 | -1.133 | -2.010 | -0.808 |
| BD70-4038 | -3.770 | -0.360 | -1.765 | -2.786 | -2.631 | 0.394 | 0.620 | -1.610 | -1.476 |
| BD70-4022 | -2.882 | -1.125 | -1.642 | -2.523 | -1.154 | 0.320 | -0.027 | -0.087 | -1.553 |
| BD70-4039 | -3.194 | -1.055 | -2.785 | -2.122 | -2.098 | -0.114 | 0.360 | -0.980 | -0.765 |
| BD70-4046 | -3.091 | -1.882 | -2.044 | -2.442 | -2.236 | 0.117 | >1 | -0.744 | -1.115 |
| BD70-4005 | -1.847 | 0.931 | -1.666 | -1.734 | -0.965 | -3.856 | >1 | 0.931 | 0.584 |
| BD70-4004 | >1 | >1 | >1 | >1 | >1 | -1.276 | >1 | >1 | >1 |
| BD70-4036 | >1 | >1 | >1 | >1 | >1 | 0.081 | >1 | >1 | >1 |
| BD70-4054 | >1 | >1 | >1 | 0.962 | 0.941 | 0.298 | >1 | >1 | >1 |
| N1D10 | >1 | >1 | >1 | 0.536 | >1 | >1 | >1 | >1 | >1 |
| BD70-4025 | >1 | >1 | >1 | >1 | >1 | 0.964 | >1 | >1 | >1 |
| SF5 | >1 | >1 | >1 | >1 | >1 | >1 | >1 | >1 | >1 |
| BD70-4051 | >1 | >1 | >1 | >1 | >1 | >1 | >1 | >1 | >1 |
| BD70-4035 | >1 | >1 | >1 | >1 | >1 | >1 | >1 | >1 | >1 |
| BD70-4032 | >1 | >1 | >1 | >1 | >1 | >1 | >1 | >1 | >1 |
| BD70-4031 | >1 | >1 | >1 | >1 | >1 | >1 | >1 | >1 | >1 |
| BD70-4029 | >1 | >1 | >1 | >1 | >1 | >1 | >1 | >1 | >1 |
| BD70-4027 | >1 | >1 | >1 | >1 | >1 | >1 | >1 | >1 | >1 |
| BD70-4015 | >1 | >1 | >1 | >1 | >1 | >1 | >1 | >1 | >1 |
| BD70-4016 | >1 | >1 | >1 | >1 | >1 | >1 | >1 | >1 | >1 |
| BD70-4023 | 0.765 | >1 | >1 | >1 | >1 | >1 | >1 | >1 | >1 |
| S2A5 | 0.471 | 0.662 | 0.327 | 0.472 | 0.412 | >1 | >1 | 0.795 | >1 |
| B1G11 | 0.447 | 0.792 | 0.839 | 0.212 | 0.973 | >1 | >1 | 1.000 | >1 |
| BD70-4017 | 0.690 | 0.132 | -1.048 | -0.842 | -0.281 | 0.831 | -1.393 | >1 | 0.202 |
| SF1 | -0.995 | 0.274 | -1.645 | -1.849 | -1.228 | 0.081 | >1 | -0.756 | -0.014 |
| BD70-4012 | -1.635 | -1.200 | -1.545 | -1.648 | -1.243 | -0.453 | >1 | -0.123 | -1.032 |
| BD70-4033 | -1.727 | -0.426 | -1.077 | -1.213 | -1.153 | 0.019 | 0.993 | -0.357 | -0.515 |
| 40C10 | -2.088 | -0.915 | -0.912 | -1.573 | -1.537 | 0.288 | 0.540 | -0.364 | -0.261 |
| BD70-4050 | -1.596 | -0.567 | -1.354 | -0.829 | -0.688 | 0.102 | -0.210 | -0.426 | -0.049 |
| BD70-4053 | -2.418 | -0.214 | -1.376 | -0.992 | -0.614 | >1 | 0.150 | 0.315 | 0.197 |

| | | | | | | | | | |
|---|---|---|---|---|---|---|---|---|---|
| BD70-4044 | -2.984 | -1.153 | -1.382 | -1.935 | -1.241 | 0.113 | 0.603 | -0.337 | 0.066 |
| BD70-4030 | -3.548 | -0.646 | -1.160 | -1.835 | -1.012 | 0.274 | 0.073 | 0.200 | 0.037 |
| BD70-4047 | -3.215 | -0.659 | -0.898 | -1.297 | -1.042 | 0.259 | 0.372 | -0.379 | -0.025 |
| BD70-4010 | -2.942 | -0.931 | -1.443 | -1.014 | -0.565 | -0.352 | >1 | 0.536 | 0.129 |
| BD70-4048 | -2.525 | -0.403 | -1.609 | -1.549 | -0.597 | 0.462 | >1 | 0.861 | 0.448 |
| BD70-4006 | -0.681 | 0.638 | -0.987 | -0.485 | -0.283 | -1.242 | >1 | 0.446 | 0.409 |
| BD70-4011 | -0.486 | >1 | -0.254 | -0.094 | -0.126 | -0.646 | >1 | 0.425 | 0.029 |
| BD70-4028 | -0.673 | >1 | -0.063 | -0.061 | 0.406 | 0.705 | >1 | 0.294 | >1 |
| bsAb3 | -0.950 | 0.143 | -0.629 | 0.005 | 0.314 | >1 | 0.521 | 0.440 | 0.796 |
| BD70-4019 | -0.337 | 0.274 | -0.005 | 0.071 | 0.247 | >1 | 0.844 | 0.574 | 0.608 |
| MAb4-5 | -0.594 | 0.539 | -0.391 | 0.178 | 0.387 | >1 | >1 | 0.805 | 0.932 |
| BD70-401 | -0.495 | 0.333 | -1.644 | -0.334 | 0.131 | 0.187 | >1 | >1 | 0.633 |
| BD70-4014 | -2.127 | 0.665 | -1.615 | -0.853 | -0.055 | 0.160 | 0.835 | 0.451 | -0.243 |
| BD70-4052 | -2.041 | -0.009 | -0.553 | -0.461 | -0.254 | 0.060 | >1 | 0.272 | 0.714 |
| BD70-4041 | -1.395 | 0.266 | -0.721 | -0.351 | 0.334 | 0.685 | >1 | 0.303 | 0.573 |
| BD70-4049 | -1.359 | 0.346 | -1.048 | -0.095 | 0.126 | 0.276 | >1 | 0.086 | 0.131 |

**Supplementary Table 5. Source data of the heatmap in Figure 1D**

| Lg IC50 (μg/ml) | HB29 | HBMC16 | AHL | SD4 | SPL030A | WCH | HN20 | G/K/2012 | HN13 |
|---|---|---|---|---|---|---|---|---|---|
| BD70-1031 | -0.870 | >1 | 0.349 | >1 | >1 | 0.356 | 0.304 | 0.354 | 0.312 |
| BD70-1009 | -0.623 | >1 | >1 | >1 | >1 | >1 | 0.570 | 0.818 | 0.533 |
| BD70-1028 | -0.559 | >1 | >1 | >1 | >1 | 0.753 | 0.684 | 0.324 | 0.441 |
| BD70-1040 | -0.572 | 0.296 | -0.165 | 0.533 | 0.518 | 0.189 | 0.019 | 0.139 | 0.069 |
| BD70-1039 | -0.545 | 0.000 | >1 | 0.727 | 0.652 | 0.288 | 0.144 | 0.213 | 0.189 |
| BD70-1034 | -0.252 | 0.227 | 0.546 | >1 | 0.901 | 0.174 | 0.664 | 0.575 | 0.766 |
| BD70-1035 | -0.328 | 0.297 | 0.099 | 0.801 | 0.748 | 0.440 | 0.513 | 0.703 | 0.625 |
| BD70-1041 | -0.230 | 0.043 | 0.195 | 0.948 | 0.647 | 0.413 | 0.540 | 0.690 | 0.685 |
| BD70-1008 | -0.213 | >1 | >1 | 0.840 | >1 | >1 | >1 | >1 | >1 |
| BD70-1025 | 0.125 | >1 | >1 | >1 | >1 | >1 | >1 | >1 | >1 |
| BD70-1042 | 0.309 | >1 | >1 | >1 | >1 | >1 | >1 | >1 | >1 |
| BD70-1037 | >1 | 0.994 | 0.128 | >1 | >1 | >1 | >1 | >1 | >1 |
| BD70-1027 | 0.879 | >1 | 0.797 | >1 | >1 | >1 | >1 | >1 | 0.959 |
| BD70-1004 | 0.872 | >1 | >1 | >1 | >1 | >1 | >1 | 0.849 | >1 |
| BD70-1014 | >1 | >1 | >1 | >1 | >1 | >1 | >1 | >1 | >1 |
| BD70-1007 | 0.531 | -0.231 | -0.099 | >1 | >1 | 0.506 | >1 | >1 | >1 |
| BD70-1010 | 0.738 | -0.027 | -0.094 | >1 | >1 | >1 | >1 | >1 | >1 |
| BD70-1002 | 0.084 | -0.553 | 0.422 | >1 | 0.552 | >1 | >1 | 0.869 | >1 |
| BD70-1029 | 0.217 | 0.248 | >1 | >1 | >1 | >1 | 0.825 | >1 | >1 |
| BD70-1030 | 0.207 | 0.184 | 0.988 | >1 | >1 | 0.682 | 0.627 | 0.876 | 0.891 |
| BD70-1001 | 0.600 | -0.231 | 0.539 | >1 | >1 | >1 | >1 | >1 | >1 |
| BD70-1003 | 0.305 | -0.304 | >1 | 0.946 | 0.884 | >1 | >1 | >1 | >1 |
| BD70-1036 | -2.613 | -1.117 | 0.506 | -0.478 | -0.899 | -0.579 | -1.327 | -1.296 | -1.019 |
| BD70-1043 | -1.579 | 0.352 | 0.990 | 0.739 | 0.594 | 0.141 | -0.288 | -0.506 | -0.360 |
| BD70-1044 | -1.332 | -0.623 | 0.454 | 0.122 | 0.249 | -0.343 | -0.704 | -0.448 | -0.260 |
| SF83 | -1.281 | -0.039 | >1 | 0.280 | 0.565 | 0.139 | 0.040 | 0.048 | -0.031 |
| BD70-1026 | -0.847 | -0.028 | 0.912 | 0.076 | 0.258 | -0.030 | -0.205 | -0.285 | -0.117 |
| BD70-1038 | -1.136 | -0.238 | 0.768 | 0.135 | -0.026 | -0.040 | -0.271 | -0.259 | -0.198 |
| BD70-1016 | -1.558 | -1.365 | -1.208 | -0.496 | -0.286 | -0.866 | -0.763 | -0.856 | -0.765 |
| BD70-1006 | -1.109 | -0.635 | -0.590 | -0.243 | 0.155 | -0.419 | 0.140 | -0.146 | -0.085 |
| BD70-1011 | -1.184 | -0.847 | -0.035 | -0.045 | -0.187 | -0.393 | -0.133 | -0.072 | -0.254 |
| BD70-1012 | -1.844 | -1.353 | -0.166 | -0.791 | -0.625 | -0.833 | -0.702 | -0.723 | -0.948 |
| BD70-1013 | -1.482 | -0.878 | -0.052 | -0.487 | -0.522 | -0.575 | -0.463 | -0.375 | -0.519 |
| BD70-1032 | -1.713 | -0.790 | 0.030 | -0.049 | -0.222 | -0.723 | -0.634 | -0.569 | -0.581 |
| BD70-1015 | -1.697 | -0.660 | -0.421 | -0.754 | -0.097 | -0.879 | -0.597 | -0.667 | -0.662 |
| BD70-1033 | -1.748 | -0.878 | -0.446 | -0.495 | 0.094 | -0.518 | -0.308 | -0.298 | -0.315 |

**Supplementary Table 6. Germline and CDR3 sequences of SFTSV-Gn or Gc antibodies**

| Antibody ID | Information of antibody sequence | | | | | |
|---|---|---|---|---|---|---|
| | Heavy chain V gene | Heavy chain J gene | Light chain V gene | Light chain J gene | CDRH3 | CDRL3 |
| BD70-4001 | IGHV3-30-3 | IGHJ6 | IGLV3-1 | IGLJ3 | AGMSALVDANYYYYGMDV | QAWDMNTAL |
| BD70-4002 | IGHV3-48 | IGHJ4 | IGLV1-44 | IGLJ3 | ELEMPTIYSPADY | AAWDDSLNGWV |
| BD70-4003 | IGHV5-51 | IGHJ6 | IGKV1-39 | IGKJ2 | HEDNGDYLTPYYVLDV | QQSFRTPST |
| BD70-4004 | IGHV3-30-3 | IGHJ4 | IGKV1-6 | IGKJ5 | FLLKYCTAGNCHSFDH | LQDYNYPIT |
| BD70-4005 | IGHV4-59 | IGHJ1 | IGKV1-5 | IGKJ1 | GWQWDDGLTNYHPLFQH | QQYNGFSMT |
| BD70-4006 | IGHV4-39 | IGHJ6 | IGLV3-21 | IGLJ1 | RRRGSSYWPTHDYYYGMDV | QVWDSGTDHLV |
| BD70-4007 | IGHV3-23 | IGHJ3 | IGLV3-21 | IGLJ2 | ELGSSTWYEPDAFEI | QVWDSSRDHWV |
| BD70-4008 | IGHV3-21 | IGHJ5 | IGLV3-21 | IGLJ3 | DPASMLGDIGELNH | QIWDTITSHRV |
| BD70-4009 | IGHV1-69-2 | IGHJ4 | IGKV3-20 | IGKJ1 | FWYFEMEPL | QQYSHSRT |
| BD70-4010 | IGHV3-49 | IGHJ4 | IGKV2-29 | IGKJ1 | SYYDFRSGRDSTTGREDY | MQGLHFPWT |
| BD70-4011 | IGHV3-15 | IGHJ4 | IGLV3-19 | IGLJ3 | GFIAVAADIDY | CSRDSSGHHLV |
| BD70-4012 | IGHV3-9 | IGHJ4 | IGLV2-14 | IGLJ3 | DISGSGGMSVAGGDY | ASFTDTNTPWV |
| BD70-4013 | IGHV1-24 | IGHJ5 | IGKV3-15 | IGKJ4 | GPYCNSANCLGWFDP | QQYNNWLT |
| BD70-4014 | IGHV1-18 | IGHJ4 | IGKV3-20 | IGKJ4 | DVGLQVVWTDYYSGSGDYYPYFLY | QQYGSSPLT |
| BD70-4015 | IGHV4-34 | IGHJ3 | IGKV3-20 | IGKJ5 | GRNPRGSQFFDWLSSGAFDI | QQYSSLHFT |
| BD70-4016 | IGHV3-33 | IGHJ3 | IGLV3-21 | IGLJ2 | DPITAVTGQDPSDALDV | QVWDSSNEHRVI |
| BD70-4017 | IGHV3-7 | IGHJ4 | IGLV1-51 | IGLJ3 | LRGWCYEGVCWGGFDL | GTWDSTLNAGV |
| BD70-4018 | IGHV3-11 | IGHJ5 | IGKV4-1 | IGKJ1 | HGNGGHYDL | QQYYSSPRA |
| BD70-4019 | IGHV4-31 | IGHJ4 | IGLV2-14 | IGLJ1 | ARINMMVVEPPPHFDY | SSYTSSSTLGV |
| BD70-4020 | IGHV4-59 | IGHJ4 | IGLV2-11 | IGLJ3 | HGSGTYPLFQFDY | CSYAGTYTWV |

| | | | | | | |
|---|---|---|---|---|---|---|
| BD70-4021 | IGHV3-48 | IGHJ4 | IGKV1-27 | IGKJ4 | DFGFSSGWSLDGSGYCDY | QKYKSAPLT |
| BD70-4022 | IGHV1-24 | IGHJ4 | IGKV3-20 | IGKJ5 | MGLLEVRFGSGTYYKYFDF | QQYGKPPIT |
| BD70-4023 | IGHV5-51 | IGHJ3 | IGLV1-44 | IGLJ3 | GRGLRGEYDI | AAWDDSLDGPGV |
| BD70-4024 | IGHV3-33 | IGHJ4 | IGLV3-21 | IGLJ2 | SRGRGYTGYDYFDL | QVWYRVGSGVQPRVV |
| BD70-4025 | IGHV3-7 | IGHJ6 | IGLV2-14 | IGLJ2 | RRPAVVVWYMDV | SSHRSSSTPVL |
| BD70-4026 | IGHV2-5 | IGHJ1 | IGKV4-1 | IGKJ2 | DYDGLYYS | QQFYGTPHT |
| BD70-4027 | IGHV3-15 | IGHJ6 | IGLV3-19 | IGLJ2 | DGEGDTVVVPAAHKYYYYYYGMDV | DSRDNTGNHLG |
| BD70-4028 | IGHV4-4 | IGHJ6 | IGLV1-44 | IGLJ3 | AGVGYRYGPSAYYYYNIDV | AAWDDILEGWV |
| BD70-4029 | IGHV3-7 | IGHJ4 | IGKV1-39 | IGKJ2 | NQWLRPFYYFDY | QQSFSAPRYT |
| BD70-4030 | IGHV3-49 | IGHJ4 | IGKV2-29 | IGKJ1 | SYYDFRSGRDLTTGREEY | MQGLHLPWT |
| BD70-4031 | IGHV3-30 | IGHJ4 | IGKV1-9 | IGKJ1 | PFRKYDVLTGCPDF | QQLSSYPWT |
| BD70-4032 | IGHV1-18 | IGHJ4 | IGLV3-1 | IGLJ2 | DSPLAATGTFLI | QGWDRSAADVL |
| BD70-4033 | IGHV2-5 | IGHJ4 | IGKV1-6 | IGKJ1 | RYYDILTGYSGLFDS | LQDYDYPWT |
| BD70-4034 | IGHV5-51 | IGHJ4 | IGKV1-39 | IGKJ1 | LLEYGDYANFIGGFDY | QYSYLVLTWT |
| BD70-4035 | IGHV3-23 | IGHJ4 | IGKV3-15 | IGKJ1 | TAGRFDY | QQYNDWPRT |
| BD70-4036 | IGHV3-9 | IGHJ6 | IGKV1-9 | IGKJ4 | PKYDHWSASQYYYLDI | QQLDSSPLT |
| BD70-4037 | IGHV3-30 | IGHJ6 | IGLV1-47 | IGLJ1 | SLAAAGTLIGYHYYGMDV | AGWDENLSTFV |
| BD70-4038 | IGHV3-30 | IGHJ3 | IGKV1-39 | IGKJ2 | TQFGELDAFDI | QQSYSTSVT |
| BD70-4039 | IGHV4-34 | IGHJ5 | IGLV1-51 | IGLJ1 | GLINFDSTGYHVRFDP | GTWDSSLGAGRGV |
| BD70-4040 | IGHV4-38-2 | IGHJ5 | IGLV1-40 | IGLJ3 | ERTSESAGMVPAEFDS | QSYDSSMSGFWV |
| BD70-4041 | IGHV3-30 | IGHJ5 | IGKV1-27 | IGKJ2 | DISAWYTGWFDP | QKYNSVPLT |
| BD70-4042 | IGHV3-23 | IGHJ4 | IGKV2-30 | IGKJ3 | SYYDYIWGTQGTDN | MQGTYRPFT |
| BD70-4043 | IGHV3-33 | IGHJ4 | IGKV1-39 | IGKJ3 | DQTDVGWNYLDS | QQSYTAPV |
| BD70-4044 | IGHV3-33 | IGHJ5 | IGKV3-15 | IGKJ4 | DIAAAATGWFDP | QQYIDWPLT |
| BD70-4045 | IGHV3-30 | IGHJ4 | IGKV3-15 | IGKJ2 | DTDYYGSGTPDV | QQYNNWPYT |
| BD70-4046 | IGHV4-34 | IGHJ4 | IGKV1-5 | IGKJ2 | GYPYHYSESGAFQSHLDS | QQHDSYPYT |

| | | | | | | |
|---|---|---|---|---|---|---|
| BD70-4047 | IGHV5-51 | IGHJ5 | IGKV3-15 | IGKJ4 | RAPGIPASPDPAFHWFDP | HQYNNGHT |
| BD70-4048 | IGHV3-74 | IGHJ6 | IGLV3-25 | IGLJ1 | GGYGFGSFYRGQENYYGMDV | QSADSSGTYV |
| BD70-4049 | IGHV5-51 | IGHJ4 | IGKV1-12 | IGKJ5 | TGRARLGRIESPAWKPFDF | QQANSFPIT |
| BD70-4050 | IGHV1-3 | IGHJ4 | IGLV1-47 | IGLJ3 | APLGPDF | AAWDDSLSGPWV |
| BD70-4051 | IGHV3-30 | IGHJ4 | IGKV2-40 | IGKJ5 | APYFDVLADYFIVDRLDQ | MQRVELPVT |
| BD70-4052 | IGHV5-51 | IGHJ4 | IGKV2D-29 | IGKJ1 | MKVRGMGAPPAFDY | MQNVHLWT |
| BD70-4053 | IGHV1-46 | IGHJ2 | IGKV1-39 | IGKJ2 | ADRGPLTPHWFFDL | HQSASAPYT |
| BD70-4054 | IGHV3-9 | IGHJ4 | IGKV3-20 | IGKJ5 | DADILYYGSRTYTPVYFDS | QQYDSSPLT |
| BD70-1001 | IGHV1-46 | IGHJ4 | IGKV3-20 | IGKJ1 | GLIRLHCGGGDCSSPLLSF | QQYGSSPRT |
| BD70-1002 | IGHV3-21 | IGHJ3 | IGLV2-14 | IGLJ1 | NDVLTGPTPKTAYREIGAFDV | SSYSTSTTLYV |
| BD70-1003 | IGHV3-30 | IGHJ4 | IGKV1-5 | IGKJ1 | DLTDTSGRLGPFDY | QQYNSYWT |
| BD70-1004 | IGHV3-9 | IGHJ5 | IGKV1-33 | IGKJ4 | HIT | QHYDTVPSLT |
| BD70-1005 | IGHV3-7 | IGHJ5 | IGLV1-44 | IGLJ3 | DSASAPDH | AAWDDGLNGWV |
| BD70-1006 | IGHV1-69 | IGHJ6 | IGKV3-20 | IGKJ2 | DDTARGTRDYYSGLDV | QQYGASPQMHT |
| BD70-1007 | IGHV1-18 | IGHJ3 | IGKV1-33 | IGKJ5 | DVSSSWSRKGHAFDI | QQYDNLPPIT |
| BD70-1008 | IGHV3-21 | IGHJ4 | IGKV3-11 | IGKJ5 | DQPATVVTPYFDY | QQRYNWPPST |
| BD70-1009 | IGHV3-21 | IGHJ5 | IGLV1-44 | IGLJ3 | LGRLSPTSFDL | SAWVDSLNGPV |
| BD70-1010 | IGHV4-39 | IGHJ3 | IGKV1-5 | IGKJ1 | HRPSDNWRNEAFDI | QQSSSYWT |
| BD70-1011 | IGHV3-30 | IGHJ4 | IGKV1-17 | IGKJ1 | ATGLITAIDS | LQYNTYPWT |
| BD70-1012 | IGHV3-48 | IGHJ4 | IGLV3-21 | IGLJ3 | KNPFEY | QVWDDNSDHRWV |
| BD70-1013 | IGHV3-30 | IGHJ4 | IGKV1-17 | IGKJ1 | ATGLITAIDS | LQYNTYPWT |
| BD70-1014 | IGHV5-51 | IGHJ6 | IGKV1-39 | IGKJ4 | DFSGTSGYPPKWFYYGMDV | QQSYSPPLT |
| BD70-1015 | IGHV3-23 | IGHJ4 | IGKV1-39 | IGKJ4 | DRAVSGWIASDS | QQSYTTPLT |
| BD70-1016 | IGHV4-59 | IGHJ4 | IGKV1-5 | IGKJ1 | GGLYDAGGYYFGY | QQYYSYRT |
| BD70-1017 | IGHV1-18 | IGHJ3 | IGKV1-39 | IGKJ4 | ELRFGELTTFDV | QQSYSSRS |
| BD70-1018 | IGHV3-74 | IGHJ5 | IGLV1-44 | IGLJ3 | DYWDLLTWVNYFDP | AAWDDSLNAWL |

| | | | | | | |
|---|---|---|---|---|---|---|
| BD70-1019 | IGHV4-39 | IGHJ3 | IGLV1-47 | IGLJ3 | RPWGYYLRDAFDI | ATWDDSLSGWV |
| BD70-1020 | IGHV3-30 | IGHJ2 | IGKV3D-15 | IGKJ1 | ERVDDYGDYATYWYFDL | QQYNNWPPWT |
| BD70-1021 | IGHV3-66 | IGHJ6 | IGKV3-20 | IGKJ2 | EPGLRNGMDV | QHFGDSPPDT |
| BD70-1022 | IGHV4-31 | IGHJ2 | IGLV1-47 | IGLJ2 | GRLDDDGVIFQWNYWYLDL | ATWDDSLNTLV |
| BD70-1023 | IGHV4-59 | IGHJ5 | IGLV3-1 | IGLJ3 | RRGSPADGFWFDP | QAWDSRFWV |
| BD70-1024 | IGHV1-18 | IGHJ4 | IGKV1-39 | IGKJ3 | ALSGSYYDYFDY | QQSYSTPFT |
| BD70-1025 | IGHV4-39 | IGHJ6 | IGLV3-25 | IGLJ3 | MEWDRIYGLDV | QSADNSGTYWV |
| BD70-1026 | IGHV4-30-2 | IGHJ4 | IGKV3-20 | IGKJ2 | DRAGAIFF | QQYGASPQMFT |
| BD70-1027 | IGHV1-3 | IGHJ3 | IGLV3-25 | IGLJ2 | KITMSTADAFDI | QSGDSSGTYVV |
| BD70-1028 | IGHV4-31 | IGHJ4 | IGKV3-11 | IGKJ4 | GGSMVRDGGYFDH | QQRGSWPFT |
| BD70-1029 | IGHV4-31 | IGHJ4 | IGKV1-16 | IGKJ2 | ARFGELLY | QQYYSYPYT |
| BD70-1030 | IGHV1-69 | IGHJ4 | IGKV1-33 | IGKJ2 | EGSRGRAKSYYDSSGPLDS | QQYDILPFS |
| BD70-1031 | IGHV3-11 | IGHJ4 | IGKV3-15 | IGKJ4 | DISTKGDYDLALDAYFGS | QQYADWPPLT |
| BD70-1032 | IGHV1-3 | IGHJ5 | IGKV1-39 | IGKJ1 | SRGVNTISGRGDWIDP | QQSYSSPPT |
| BD70-1033 | IGHV3-7 | IGHJ6 | IGLV3-1 | IGLJ3 | GGSTGYNYGMDV | QTWDSSTLV |
| BD70-1034 | IGHV1-46 | IGHJ4 | IGKV3-20 | IGKJ4 | GVPQGGGYYSGTIDY | QQYGSSPLLS |
| BD70-1035 | IGHV3-21 | IGHJ4 | IGLV1-40 | IGLJ1 | DGYIGPAGTYLL | QSYDSSLSGSV |
| BD70-1036 | IGHV4-34 | IGHJ5 | IGKV3-15 | IGKJ2 | SGGIRVAQLLFGGPNWFDP | QQYNNWPYT |
| BD70-1037 | IGHV3-30 | IGHJ4 | IGKV1-17 | IGKJ4 | TGAKVAADDY | LQHNTYPLT |
| BD70-1038 | IGHV1-69 | IGHJ4 | IGKV1-17 | IGKJ2 | DSAWLPDY | LQHNGFPYT |
| BD70-1039 | IGHV1-46 | IGHJ6 | IGLV3-9 | IGLJ3 | DAAPLNCGGDCLYYGLDV | QVWDNTELV |
| BD70-1040 | IGHV3-23 | IGHJ4 | IGKV1D-12 | IGKJ4 | EKVVAAAGFWFDY | QQANTFPLT |
| BD70-1041 | IGHV1-18 | IGHJ2 | IGKV4-1 | IGKJ3 | QGYYDSSGYFTYWYFDL | QQYFSSPPS |
| BD70-1042 | IGHV3-33 | IGHJ6 | IGLV1-40 | IGLJ1 | GPQTEEEPRSSYYYYHGLDV | QSYDNSLNVYV |
| BD70-1043 | IGHV1-69 | IGHJ2 | IGKV3-20 | IGKJ1 | GGGIAAAGPGPPPMDL | QQYGSSPWA |
| BD70-1044 | IGHV4-61 | IGHJ4 | IGKV1-39 | IGKJ1 | VDNSGWFAIDN | QQSYITPRT |

**Supplementary Table 7. SHM counts and rates in nucleotides, in addition to the germline V-J gene combinition of the 6 antibodies.**

| Antibody | Epitope Group | SHM H (nt) | SHM L (nt) | SHM H rate (nt) | SHM L rate (nt) | Heavy chain V gene | Heavy chain J gene | Light chain V gene | Light chain J gene |
|---|---|---|---|---|---|---|---|---|---|
| BD70-4003 | IA | 27 | 30 | 0.072 | 0.0932 | IGHV5-51 | IGHJ6 | IGKV1-39 | IGKJ2 |
| BD70-4008 | IIIA | 22 | 28 | 0.0595 | 0.0862 | IGHV3-21 | IGHJ5 | IGLV3-21 | IGLJ3 |
| BD70-4009 | IIIA | 14 | 13 | 0.0394 | 0.0404 | IGHV1-69-2 | IGHJ4 | IGKV3-20 | IGKJ1 |
| BD70-4013 | IA | 9 | 8 | 0.0241 | 0.0251 | IGHV1-24 | IGHJ5 | IGKV3-15 | IGKJ4 |
| BD70-4017 | IIIA | 30 | 15 | 0.0798 | 0.0453 | IGHV3-7 | IGHJ4 | IGLV1-51 | IGLJ3 |
| BD70-4022 | IIIA | 14 | 15 | 0.0364 | 0.0462 | IGHV1-24 | IGHJ4 | IGKV3-20 | IGKJ5 |

**Supplementary Table 8. Cryo-EM data collection, refinement and validation statistics**

| | HB29-GN-head-3-17 complex Consensus map | HB29-GN-head-8-17 complex Consensus map | HB29-GN-head-3 complex Consensus map | HB29-GN-head-8 complex Consensus map |
|---|---|---|---|---|
| Data collection and processing | | | | |
| Microscope | KriosG4 | KriosG4 | KriosG4 | KriosG4 |
| Camera | Falcon 4 | Falcon 4 | Falcon 4 | Falcon 4 |
| Imaging mode | Counted resolution | Counted resolution | Counted resolution | Counted resolution |
| Magnification | 130k | 130k | 130k | 130k |
| Voltage (kV) | 300 | 300 | 300 | 300 |
| Electron exposure (e–/Å2) | 60 | 60 | 60 | 60 |
| Defocus range (μm) | -1.0~-2.0 | -1.0~-2.0 | -1.0~-2.0 | -1.0~-2.0 |
| Pixel size (Å) | 0.95 | 0.95 | 0.95 | 0.95 |
| Symmetry imposed | C1 | C1 | C1 | C1 |
| Initial particle images (no.) | 1,921,197 | 3,003,207 | 2,117,069 | 2,093,432 |
| Final particle images (no.) | 477,156 | 509,899 | 183,639 | 124,291 |
| Map resolution (Å) | 2.69 | 2.69 | 2.85 | 2.97 |
| FSC threshold | 0.143 | 0.143 | 0.143 | 0.143 |
| Map resolution range (Å) | 1.8-5.0 | 1.8-5.0 | 2.0-5.0 | 2.0-5.0 |
| Refinement | | | | |
| Initial model used (PDB code) | AF3 predicted | AF3 predicted | AF3 predicted | AF3 predicted |
| Map sharpening B factor (Å2) | -76.7 | -80.4 | -86.3 | -68.7 |
| Model composition | | | | |
| Protein residues | 788 | 787 | 546 | 546 |
| Ligands | 11 | 11 | 9 | 9 |
| R.m.s. deviations | | | | |

| | | | | |
|---|---|---|---|---|
| Bond lengths (Å) | 0.003 | 0.002 | 0.003 | 0.004 |
| Bond angles (°) | 0.681 | 0.525 | 0.547 | 0.585 |
| MolProbity score | 1.4 | 1.12 | 1.18 | 1.28 |
| Clashscore | 3.32 | 3.26 | 2.48 | 3.7 |
| Validation | | | | |
| Rotamers outliers (%) | 0 | 0 | 0.21 | 0 |
| Cβ outliers (%) | 0 | 0 | 0 | 0 |
| Ramachandran plot | | | | |
| Favored (%) | 96.02 | 98.07 | 97.21 | 97.4 |
| Allowed (%) | 3.73 | 1.93 | 2.79 | 2.6 |
| Disallowed (%) | 0.26 | 0 | 0 | 0 |
